# Harnessing Gradients for Self-Assembly of Peptide-Based Nanocapsules: A Pathway to Advanced Drug Delivery Systems

**DOI:** 10.1101/2023.07.19.549672

**Authors:** Haopeng Li, Xuliang Qian, Harini Mohanram, Xiao Han, Huitang Qi, Guijin Zou, Fenghou Yuan, Ali Miserez, Qing Yang, Tian Liu, Huajian Gao, Jing Yu

## Abstract

Biological systems often create materials with intricate structures to achieve specialized functions. In comparison, precise control of structures in man-made materials has been challenging. Here, we report a serendipitous discovery of insect cuticle peptides (ICPs) spontaneously forming nanocapsules through a single-step solvent exchange process, where the concentration gradient resulting from mixing of water and aceton drives the localization and self-assembly of the peptides into hollow nanocapsules. The underlying driving force is the intrinsic affinity of the peptides for a particular solvent concentration, while the diffusion of water and acetone creates a gradient interface that triggers peptide localization and self-assembly. This gradient-mediated self-assembly offers a transformative pathway towards next-generation drug delivery systems based on peptide nanocapsules.

## Introduction

In the natural process of evolution, organisms have developed sophisticated strategies to adapt to their harsh environments by utilizing simple raw materials and optimizing their functional properties^1^. One such strategy is the incorporation of functional gradients in the assembly of biomaterials^2^, which involve changes in local chemical composition^3-5^, arrangement of building units^6-8^, and distribution within materials^9,10^. These gradients can be observed at various length scales in biological materials. However, achieving precise control over gradients remains a formidable task in artificial self-assembly systems.

Artificial creation of such biomaterials often necessitates the engineering of a gradient interface, which can be a challenging endeavor. Particularly, solvent plays critical role in creating such gradients in numerous self-assembly systems^11-13^. By regulating the interactions between the solvent and the components of the assembly, a range of nano-sized structures can be formed, such as particles^14-16^, fibers^12,17,18^, films^19,20^, or vesicles^21,22^. These assembly processes often exhibit dynamic^23^ and non-equilibrium characteristics^24,25^. Despite their importance, the mechanisms underlying these assembly processes are often not well-understood. Nanoprecipitation, a common solvent mediated assembly technique, leverages mixed solvents to steer precipitation-based self-assembly, resulting in the production of polymer nanoparticles. The technique involves adding an excess of a poor solvent to a polymer solution, which triggers the formation of solid polymer nanoparticles through precipitation. This process has been successfully scaled up for industrial applications in fields as diverse as pharmaceuticals and agriculture^26^.

Here, we unveil a new strategy for harnessing gradients in the design of biomimetic functional self-assembling peptides. Leveraging the proteome of insect epidermis, we have identified that three selected biomimetic peptides with repetitive sequences, can form hollow nanocapsule structures without the need for any external templates. The nanocapsules formation is first initiated by the rapid formation of peptide-rich microdroplets, followed by a slower solvent gradient-mediated self-assembly process. The primary driving force for the initial phase separation is the affinity of the peptides for the water-rich environment. At a longer timescale, the diffusion between water and acetone creates a gradient interface, disrupting the metastable microdroplet state and prompting the localization and self-assembly of peptides in the vicinity of this gradient interface. These peptide-based nanocapsules can be a versatile delivery platform with potential applications in various biomedical fields, such as drug delivery, immunotherapy, and gene therapy.

## Results and discussion

### Screening of the Insect Cuticle Inspired Peptides

We conducted a label-free, quantitative proteomic analysis on the head capsule of *O. furnacalis*, (Fig. 1a and Supplementary Fig. 1). This analysis yielded a total of 951 proteins, identified through a search in the SwissProt database. The normalized spectral abundance factor (NSAF), calculated based on spectral counting, was used to quantify each protein from the proteomic data sets. A subsequent transcriptomic analysis of *O. furnacalis* larvae cuticle revealed 233 cuticle proteins derived from unigenes (Fig. 1b and Supplementary Table 1). These cuticle proteins possess molecular weights ranging from approximately 5 to 60 kDa. A comparative analysis with the amino acid compositions of eukaryote proteins revealed that cuticle proteins are heavily skewed towards Gly, Ala, and Val, with a moderate preference for Leu, Pro, and Ser^27^ (Supplementary Fig. 2). Given that functionally significant cuticle proteins often contain recurring functional domains, such as the chitin-binding Rebers and Riddiford (RR) domains^28^, we conducted a further examination of the polypeptide sequences derived from the cuticle proteins. Our screening criteria for these sequences necessitated the presence of three or more repeating units, each comprising a minimum of five amino acids. Adhering to these screening criteria, we were able to isolate nine repeated peptides from the insect cuticle protein sequences (Fig. 1d, Supplementary Fig. 3 and Supplementary Table 2).

**Fig. 1.**
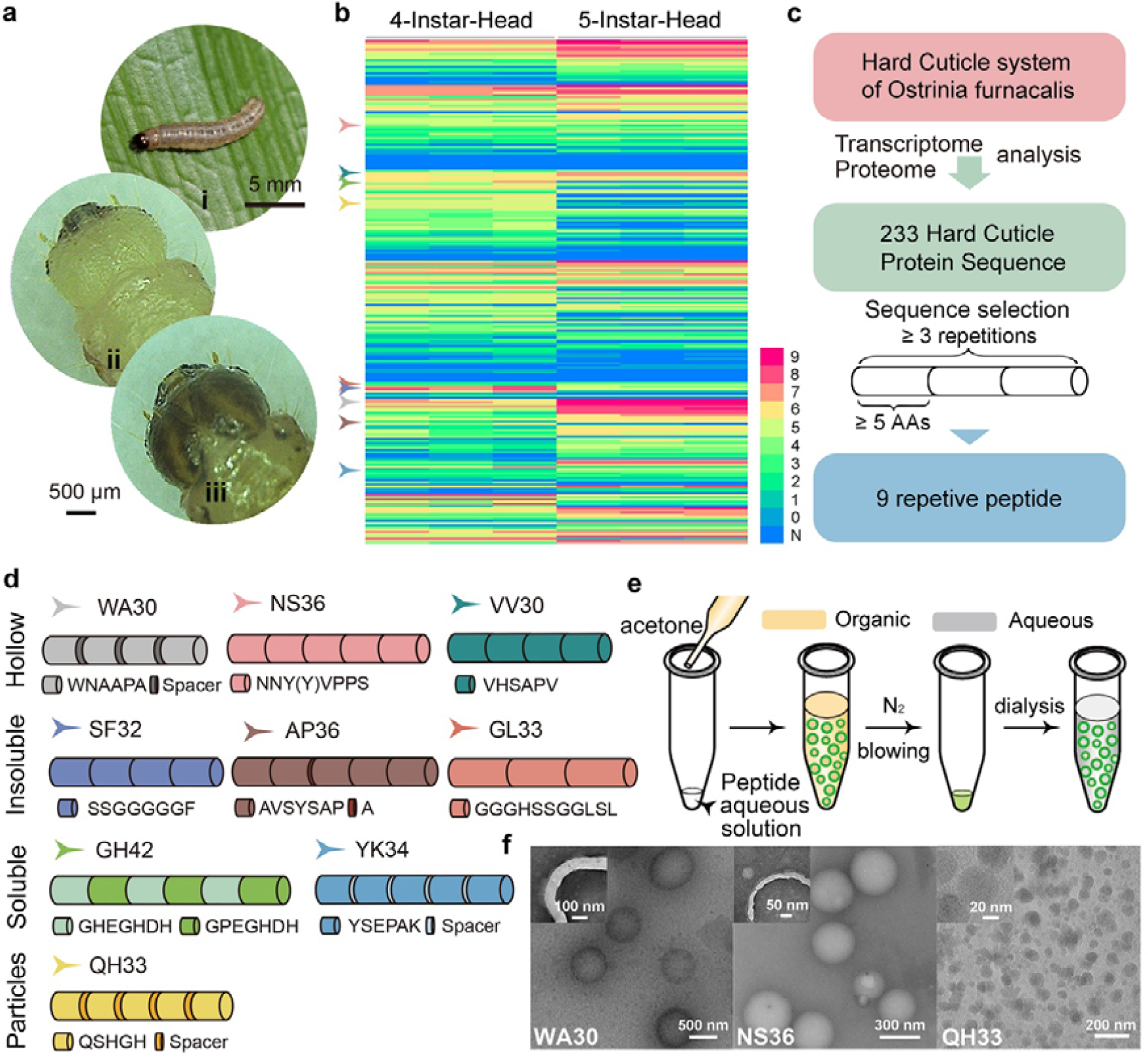
*De novo* sequencing and screening of nanocapsule forming peptides from *O. furnacalis*. **a**, Images of an *O. furnacalis* larva (i) and its head capsule 0 min (ii) and 75 mins (iii) after the fourth molting. **b**, Heat map showing the abundance of identified proteins in the *O. furnacalis* head capsule transcriptome. **c**, Screening of ICP sequences from cuticular proteins. **d**, Sequences of the selected ICPs. **e**, Schematic illustration of the peptide nanocapsule preparation process. **f**, TEM images of WA30 (left) and NS36 (middle) peptide nanocapsules, and QH33 (right) peptide nanoparticles. The insets show the enlarged nanocapsules or particles. WA30 and NS36 nanocapsules were stained with 0.3% uranium acetate solution for 15 seconds prior to imaging.

### Mixed solvent induced self-assembly of insect cuticle inspired peptides

We investigated the assembly of the insect cuticle peptides (ICPs) in mixed solvents using various ratios of water and acetone, employing the nanoprecipitation method^29^ (Fig. 1e and Methods). In this method, acetone serves as a poor solvent for the peptides, enabling the induction of peptide assembly upon the addition of multiple acetone volumes into a concentrated peptide aqueous solution. The interaction between the solvent and peptides plays a vital role in controlling peptide assembly. Non-polar solvents, such as diethyl ether^30^, result in extensive peptide precipitation, while solvents with high polarity, like dimethyl sulfoxide^31,32^, lead to no observable assembly (Supplementary Table 3). Five peptides, namely GL33, SF32, AP36, GH42, and YK34, were unable to form assembly structures due to either low solubility in water (GL33, SF32, and AP36) or excessive charge presence (GH42 and YK34). In contrast, four peptides, namely WA30, NS36, VV30, and QH33, successfully formed nanostructures. We introduced isophorone diisocyanate (IPDI) as a crosslinking agent during the assembly process, serving to crosslink the N- and C-termini of the assembled peptides^33,34^. It is noteworthy that the presence of IPDI ensures the structural integrity of the peptide assemblies in further aqueous solution applications but does not affect the assembly structure of the peptides in the mixed acetone/water solution (Fig. 1f and Supplementary Fig. 4).

Peptides NS36, WA30, and VV30 demonstrated solvent-tunable secondary structures (Supplementary Fig. 5). We hypothesized that this characteristic would enable the modulation of these peptides assemble by adjusting the peptide-solvent interactions. Intriguingly, we discovered that by altering the volume ratio of water and acetone during the assembly process, these three peptides could be coaxed to form hollow nanocapsules. As per our understanding, the generation of such hollow nanocapsule structures is a rare occurrence during the nanoprecipitation process. In contrast, peptide QH33 formed solid nanoparticles, which aligns with the expected outcome from a nanoprecipitation process.

### Assembly of ICP capsules is mediated by microdroplets

Our experimental observations reveal the presence of two distinct timescales in the self-assembly process of ICPs without the presence of crosslinker. The assembly process is initiated by forming peptide-rich microdroplets (Fig. 2a). Immediately upon the addition of acetone, we witnessed the formation of liquid droplets for NS36, WA30, and VV30. The required minimum acetone-to-water volume ratios for phase separation, using peptide saturated aqueous solutions, were 2.41, 3.65, and 3.76, respectively. By integrating microscopic observations and solution turbidity measurements, we identified the peptide concentrations and acetone-to-water volume ratios that resulted in the microdroplet formation for these three ICPs (Fig. 2b and Supplementary Fig. 6).

**Fig. 2.**
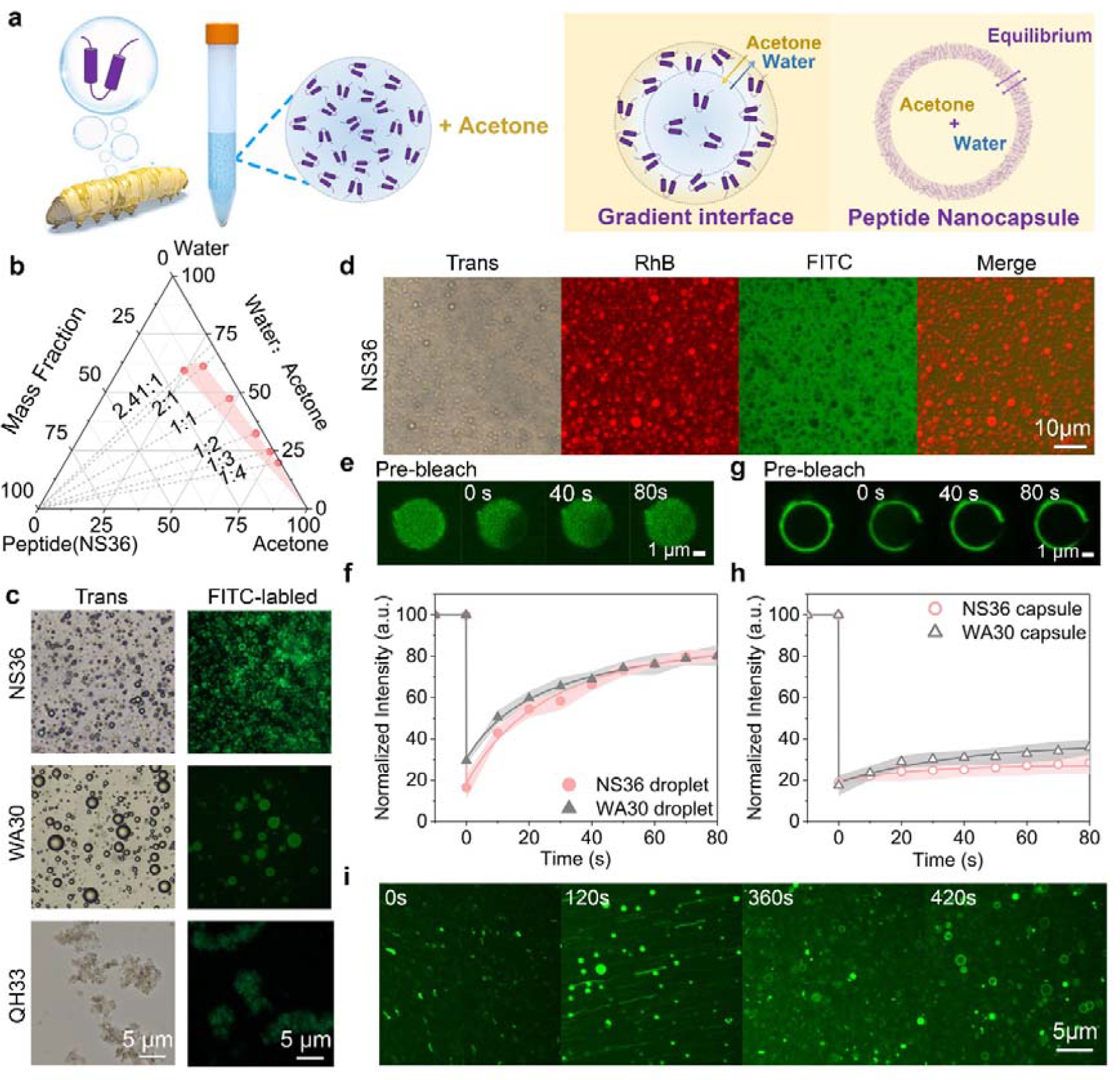
The assembly of ICPs in water-acetone mixed solvent. **a**, Schematic illustration of the experiment setup and solvent gradient-mediated assembly of ICP capsules. **b**, Ternary phase diagram showing the phase separation condition of solutions with acetone, water, and NS36 peptide. The pink area indicates the phase separation region. **c**, Optical (left) and fluorescence (right) images of peptide assemblies (2 mg/mL) formed after 10 minutes of microdroplet formation at a 9:1 volume ratio of acetone to water. **d**, Fluorescence distribution micrograph of freshly prepared NS36 droplets infiltrated with FITC and Rhodamine-B, taken after the final mixed solution was prepared under a 9:1 volume ratio of acetone to water and a final peptide concentration of 4 mg/mL. **e-f**, Representative images (e) and FRAP curve (f) of freshly prepared droplets of NS36 (2 mg/mL) under a 9:1 volume ratio of acetone to water. **g-h**, Representative images (g) and FRAP curve (h) of fully matured NS36 capsules (2 mg/mL) taken 10 minutes after microdroplet formation. Data are presented as mean values ± SD from three repeated experiments. **i**, Time-varying confocal micrograph showing the transformation of NS36 capsules from the microdroplet state.

The rapid formation of peptide-rich microdroplets upon the introduction of acetone into the peptide-water solution implies the operation of a fast timescale in the early stages of peptide assembly. Subsequently, a lengthier timescale, extending over several hundred seconds, is needed to transition the peptide droplets to more solid-like structures (Fig. 2e-h). Through optical and fluorescence microscopy imaging of fluorescein isothiocyanate (FITC) labelled ICPs, we observed that the droplet size continuously increased over several hundred seconds due to droplets coalescence (Fig. 2c). Dynamic Light Scattering (DLS) measurements further confirmed this trend. The final microdroplet sizes were recorded as 0.9 ± 0.14, 2.3 ± 0.28, and 4.2 ± 0.87 μm for VV30, NS36, and WA 30, respectively. The size of the droplets can be tuned by the amount of IPDI added. (Supplementary Fig. 7).

The swift formation of microdroplets Is facilitated by the ICPs, given that water and acetone can be combined in any ratio. Acetone acted as a poor solvent for the peptides, leading us to speculate that the peptide-rich droplets are primarily water-based. This hypothesis was validated using fluorescent dyes with varying degrees of hydrophobicity. The water-soluble dye, Rhodamine-B (Rh-B)^35^, was found to accumulate in the microdroplets, whereas the more hydrophobic Fluorescein isothiocyanate (FITC) remained in the surrounding solution^36^ (Fig. 2d and Supplementary Fig. 8). In essence, such phase separation results in a gradual transition zone rather than a sharply defined interface between the two phases. This zone is characterized by a steady evolution of acetone concentration, maintaining constant water-acetone diffusion as long as a concentration gradient exists^37^ (Fig. 2a).

Fluorescence microscopy observations indicated that the ICP microdroplets transitioned from homogeneous liquid droplets into capsules with hollow centers approximately 6 minutes after the initial mixing (Fig. 2e and Supplementary Fig. 9). To gain further insight into this microdroplet mediated peptide self-assembly process, we carried out fluorescence recovery after photobleaching (FRAP) experiments using Alexa 488 labeled NS36 and WA30 (Methods, Fig. 2e-g and Supplementary Fig. 10). However, the VV30 droplets were not suitable for FRAP measurement due to their exceedingly small diameter. Following photobleaching, both NS36 and WA30 droplets exhibited nearly 80% fluorescence recovery in 80 seconds, indicating the fluidic state of the microdroplets (Fig. 2f and Supplementary Fig. 11). In contrast, after the formation of capsules, the fluorescence intensity of the capsule walls in NS36 and WA30 only recovered by 27.6% and 37.8%, respectively (Fig. 2h). This suggests that during the peptide capsule assembly process, the peptides in the droplets undergo a transition from a liquid state to a gel state^38,39^.

### β-sheet formation promotes the assembly of ICPs

To probe the phase separation mediated ICP assembly mechanism, we examined the ICP secondary structures during assembly using circular dichroism (CD) and ATR-FTIR. Capsule-forming peptides WA 30, VV30, and NS36, exhibited distinct solution structures in water: WA 30 displayed α-helical characteristics, while VV30 and NS36 revealed a mix of extended coil and β-turn structures. The nanoparticle forming QH33 showed a random coil structure^40^ (Fig. 3a and Supplementary Fig. 12). Further 2D ^1^H-^1^H TOCSY and ^1^H-^1^H NOESY measurements confirmed these observations. ^1^H-^1^H NOESY and 2D ^1^H-^15^N HSQC spectra revealed that WA30 mainly contained helical conformation, evidenced by the i, i+3 and i, i+4 NOEs spanning over residues S16-A28, while QH33 mainly exhibited an extended conformation^41^. 2D ^1^H-^15^N HSQC spectra of NS36 suggested a flower like loop structure (Fig. 3b,c and Supplementary Fig. 13-16).

**Fig. 3.**
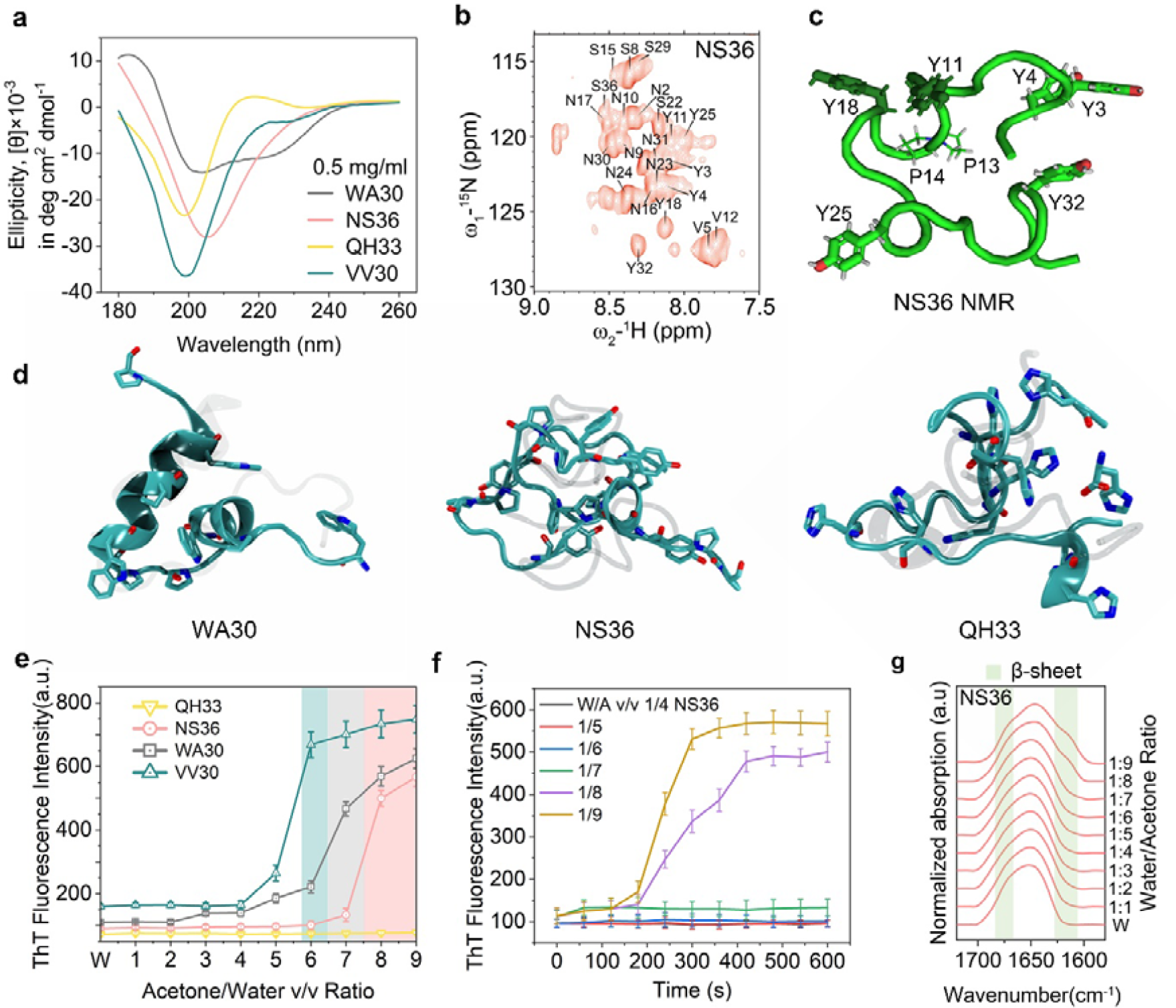
Transition of ICP secondary structure during the self-assembly process. **a**, CD spectra of ICPs (0.5 mg/mL) in aqueous solutions. **b**, 2D ^1^H-^15^N HSQC spectrum of NS36 with individual amino acids labeled. **c**, Single representative energy-minimized structure of NS36 from NMR measurement. **d**, Simulation snapshots of single peptide WA30 (left), NS36 (middle), and QH33 (right) in water. The backbone atoms of each peptide were aligned to the corresponding NMR structures (transparent) using VMD RMSD calculator (WA30: 3.17 Å; NS36: 6.96 Å; QH33: 3.36 Å). **e**, Fluorescence spectra of ICP transformation structure information in gradient-composed mixed solvents monitored through ThT staining. **f**, Kinetic analysis of β-sheet evolution during capsules formation based on ThT staining monitoring; **g**, The amide I ATR-FTIR spectra of NS36 (2 mg/mL) under different water-to-acetone volume ratios. All data are presented as means ± SD binding assay from n = 3 independent assays.

We also corroborated the solution conformations of ICP through atomistic simulations, using structures obtained from liquid-phase NMR experiments as initial configurations for aqueous MD simulations. The simulated peptide configurations showed stable root-mean-square deviation (RMSD), solvent accessible surface areas (SASA), and radius of gyration (Rg) evolution, suggesting peptide stability within the simulation timescale in water, akin to the NMR experiments. The obtained configurations from MD exhibited a high degree of resemblance to the NMR structures, thereby validating the efficacy of MD simulations as a valuable tool for investigating the behavior of ICP (Fig. 3d and Supplementary Fig. 17,18).

Upon forming peptide capsules, the ICPs transition into β-sheet structures^42^. This is supported by the strong fluorescence displayed by the ICP capsules upon staining with thioflavin-T (ThT), which indicates the presence of cross-β-sheet structures in the capsule walls (Supplementary Fig. 19). To monitor the assembly kinetics of the ICPs within the microdroplets, we conducted a ThT assay to track fluorescence emission based on water to acetone ratio and time. No capsule formation was observed for NS36 at water content (vW/vA) higher than 1:7, and fluorescence emission remained stable over time (Fig. 3e). In conditions leading to droplet formation (1/9 ≤ vW/vA ≤ 1/8), weak fluorescence was detected in the initial 150 seconds, then rapidly escalated, plateauing within 100-200 seconds -coinciding with the formation of peptide capsules (Fig. 3f). Thereafter, the ThT fluorescence intensity was continuously monitored for six hours and remained steady even after six months (Supplementary Table 4). WA30 and VV30 showed similar behavior, with an increase in acetone volume fraction leading to faster kinetics of β-sheet formation and higher β-sheet content (Supplementary Fig. 19). The formation of β-sheets during the capsule forming process was also confirmed by ATR-FTIR results (Fig. 3g and Supplementary Fig. 20-23).

### Uncovering the driving forces of ICPs self-assembly

During the introduction of acetone into the peptide-water solution, a rapid phase separation occurs, resulting in the formation of water-rich droplets containing peptide molecules, surrounded by an acetone-rich phase. This process does not create a sharp interface between the two phases, but rather a gradient interface characterized by a transition zone with smoothly changing acetone concentration (Fig. 2a). Notably, simulation studies revealed the swift diffusion of acetone into the water-rich phase, with a diffusion coefficient on the order of 1 × 10^−5^ cm^2^/s (Supplementary Fig. 24 and Supplementary Table 5). This diffusion process is instrumental in defining the concentration profile and location-specific chemical environments at the gradient interface. Consequently, it prompts the vital question of how such a dynamic gradient interface could influence the self-assembly and encapsulation behavior of ICPs, thus driving our efforts to unearth the driving forces at play.

We hypothesized that peptide-peptide interactions vary with changes in the mixed solvent. To investigate these interactions and determine the timescale guiding the initial peptide assembly stages, we carried out MD simulations of numerous isolated peptides at a variety of pre-equilibrated acetone concentrations, reflecting the experimental conditions (i.e., vA/vW = 0,1,3,5,7,9). Initial contact of peptides is driven by π -π interaction between tryptophan (W) residues for WA30, hydrogen bonding networks for NS36, and stable hydrogen bond pair between the side chains of Glu and His for QH33. (Fig. 4a-c). We observed intriguing results relating to the timescale required for the formation of initial self-assemblies (Supplementary Table 6). This timescale ranged from a few tens of nanoseconds to hundreds of nanoseconds, depending on both acetone concentration and peptide sequence. Importantly, we found that higher acetone concentrations were correlated with longer times required for the initial contact of NS36, influencing the early stages of peptide self-assembly.

**Fig. 4.**
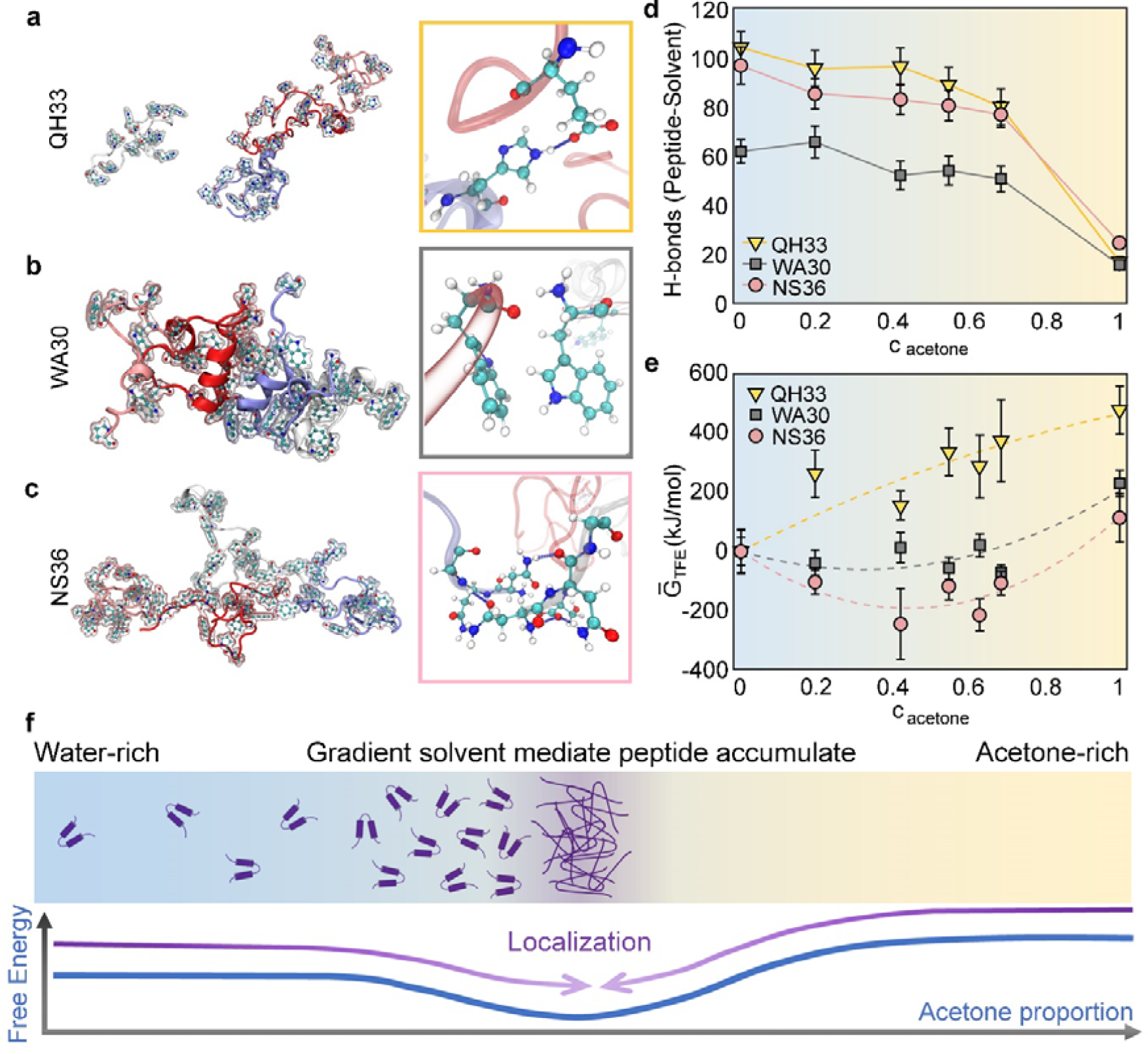
Exploring the driving forces of ICP self-assembly through Molecular Dynamics (MD) simulations. **a-c**, Simulation snapshots of the self-assembly of QH33 (a), WA30 (b) and NS36 (c), in the mixed solution with acetone concentration of 0.63. Four peptides (red, pink, silver, and blue-grey) were simulated for each case. Insets: zoom-in views of the detailed inter-peptide interactions. **d**, The number of H-bonds between a peptide and the solvent molecules (water and/or acetone) plot against the concentration of acetone. **e**, Free energy profile of transferring a peptide from pure water to the target chemical environment (c_acetone_) showing different trend for QH33 versus WA30 and NS36. The dashed lines mark the quadratic fitting of each corresponding ΔG□_TFE_ with intercepts setting to zero at c_acetone_ = 0. Data are means□±□SD calculated with the Multistate Bennett Acceptance Ratio (MBAR) method. **f**, A graphical schematic showcasing dynamic processes and driving force for gradient-mediated self-assembly of nanocapsule forming peptides.

We assessed the number of hydrogen bonds (H-bonds) between peptides and solvent at each simulated concentration. The total H-bonds between peptides and solvent decreased mildly at high acetone concentrations (*cacetone* 0.69, or vA/vW = 9) for NS36 and WA30 (by 4.35% and 1.83%, respectively), and significantly for QH33 (by 21.22%) compared to the reference concentration (*cacetone* 0.42, or vA/vW = 3). In pure acetone, the number of H-bonds decreased drastically for all three peptides (Fig. 4d and Supplementary Fig. 25). Additional analysis of SASA, RMSD, and Rg suggested that peptide assembly adopted more extended conformations as acetone concentration increased in the solvent. The exception was QH33 at concentrations above 0.42 (or vA/vW = 3), which coincided with the turning point in H-bond numbers between peptides and solvent molecules (Supplementary Fig. 26).

In view of the different timescales between experiments and MD simulations, we performed simulations on the peptide-peptide interactions in the pre-equilibrated mixed solvent, noting that the diffusion of peptides in solvents has been reported to be significantly slower, with diffusion coefficients typically one to two orders of magnitude smaller compared to water-acetone diffusion^43^. Furthermore, we conducted Free Energy Perturbation (FEP) simulations to calculate the transfer free energy (TFE) of peptides at different concentrations across the water-acetone interface (Methods and Fig. 4e).

Our calculated TFE landscape reveal a consistent and incremental increase for the nanoparticle-forming peptide QH33 as the acetone concentration rises. This trend suggests an energetic disadvantage for QH33 as acetone molecules infiltrate the peptide containing water-rich phase. The incursion of acetone alters the chemical environment surrounding the QH33 peptides, driving their migration towards a more aqueous environment with higher water concentration. This relocation ultimately culminates in the precipitation of QH33 peptides, and such process conforms to nanoprecipitation principles.

Our investigation revealed important insights into the behavior of nanocapsule forming peptides, NS36 and WA30. These peptides exhibit single-well TFE landscapes with global minima observed between pure water and pure acetone. These energy minima mark the energetically optimal concentrations for peptide localization at the forefront of water-acetone mixing. The free energy differences provide the driving force for peptides to spontaneously migrate towards the optimal concentration at the gradient interface, triggering peptide localization and facilitating the nanocapsule formation (Fig. 4f). It is worth noting, however, that not all peptides manifested this behavior—only those displaying local minima in their TFE landscapes showed the propensity for nanocapsule formation.

We underscore that these observations from molecular dynamics simulations and free energy perturbation calculations are rare but not unexpected. In fact, acetone does not act strictly as a “poor solvent” for nanocapsule forming peptides. Rather, the concentration of acetone plays a crucial role in the formation of nanocapsules. Insufficient amounts of acetone fail to create necessary conditions for capsule formation, while excessive concentrations still allow the formation process to occur (Fig. 3e). The localization of the ICPs at the water-acetone interface, coupled with a high acetone concentration locally, facilitates the formation of β-sheet structures occurring over an extended timescale. Furthermore, we propose that peptide localization at the gradient interface may impede the diffusion of acetone into the water phase. This barrier to water-acetone diffusion might account for the longer timescale (>100 seconds) for microdroplet to nanocapsule transition observed in the experiments.

### ICP capsule based cytosolic delivery of drugs

The solvent gradient-mediated assembly process and hollow structure of the ICPs make them effective vehicles for co-delivering large hydrophilic biomolecules and small molecule drugs^44^. Early in the phase separation process, hydrophilic biomacromolecules, such as nucleotides and proteins, prefer to reside in the water-rich droplet phase alongside the ICPs. During the solvent exchange process, small hydrophobic molecules dissolved in acetone can be concentrated and trapped in the capsules (Fig. 5a). We utilized enhanced Green Fluorescent Protein (EGFP) and Doxorubicin (DoX) as model compounds for large hydrophilic and small hydrophobic drugs to demonstrate this capability. All ICP capsules exhibited exceptionally high EGFP loading efficiency, ranging from 87.5% to 92.1% (Fig. 5b). Similar loading efficiencies were observed for other biomacromolecules, including DNA, RNA, β-Galactosidase, and Smac^45^, using NS36 nanocapsules (Supplementary Fig. 27). In contrast, the loading efficiency for small molecules was highly sequence-dependent; NS36 capsules displayed the highest loading efficiency (∼ 54.8%) for DoX, whereas VV30 demonstrated the highest loading efficiency (∼76.2%) for the more hydrophobic drug, paclitaxel (Fig. 5c).

**Fig. 5.**
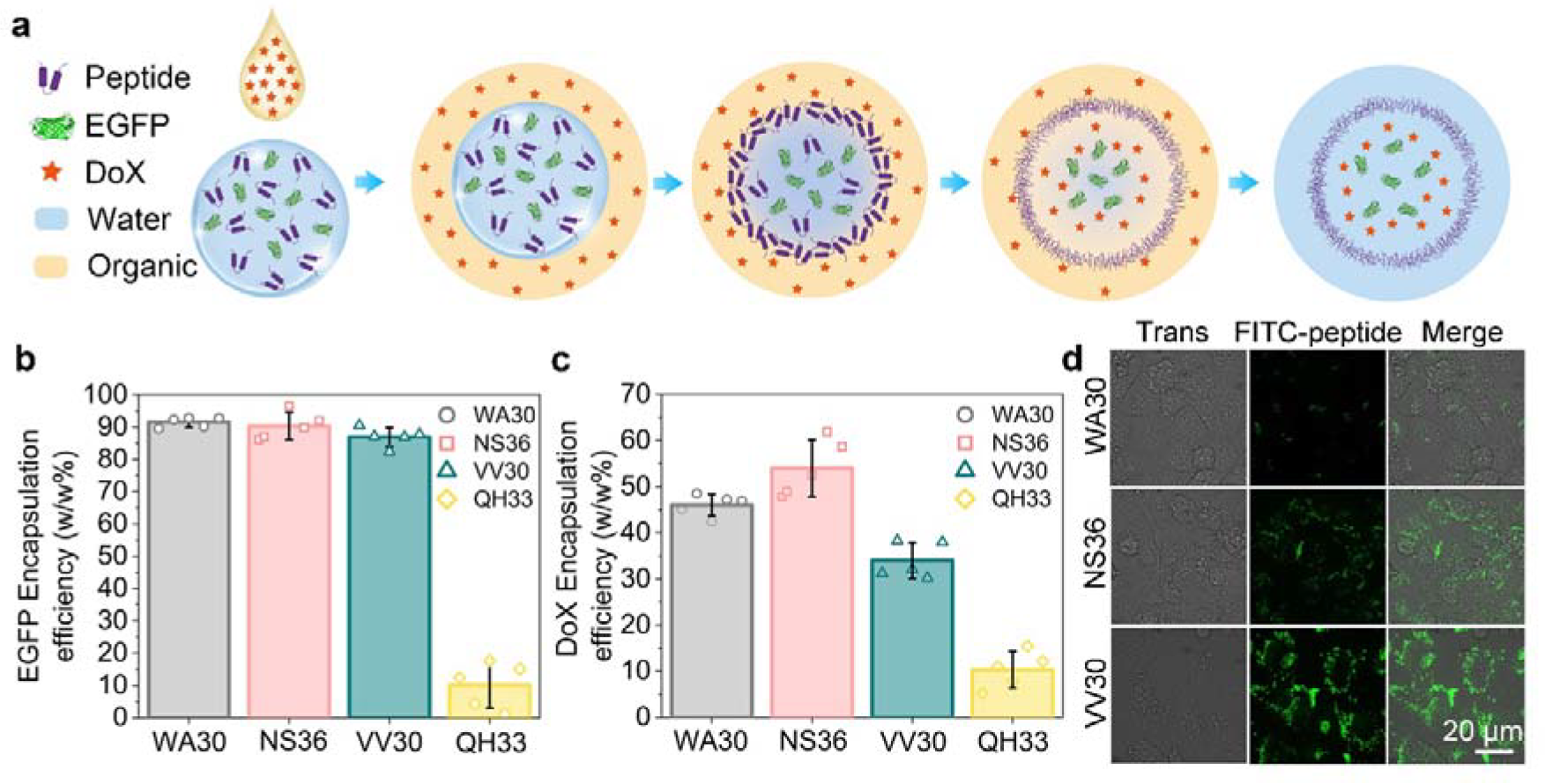
Encapsulating protein and drug by ICP capsules. **a**, Schematic representation of encapsulating DoX and EGFP simultaneously into the ICP nanocapsules. **b-c**, EGFP (b) and DoX (c) encapsulation efficiency of the ICP capsules. Data are displayed individually (hollow dots of the same color as the bar) and as the mean□±□SD (column with error bar) from n□=□5 independent measurements. **d**, Representative fluorescence images demonstrating the successful delivery of FITC-labelled ICP nanocapsules into Hela cells. All delivery results were confirmed by n□=□3 independent assays.

The cellular uptake of FITC-labeled peptide nanocapsules in Hela cells was confirmed through confocal microscopy (Fig. 5d). The ICP capsules demonstrated very low cytotoxicity, as indicated by the MTT assay, with over 90% cell survival for all three peptides even at a concentration of 400 mg/mL (Fig. 6a). Confocal microscopy images revealed that VV30 exhibited the highest uptake efficiency, while WA30 showed the lowest efficiency, likely due to its larger size^46^ (Supplementary Fig. 28). Furthermore, we conducted examinations on the uptake of NS36, a compound that was labeled in U2Os and MCF7 cells. Remarkably, our findings revealed substantial cytotoxicity along with notable uptake efficiency across both cell lines. To further investigate the versatility of the ICP capsules, the FITC-labeled NS36 capsules were tested on another two cancer cell lines, namely MCF7 and U2OS. Based on the fluorescence signals observed inside the cells, the low cytotoxicity and high uptake efficiency of NS36 capsules was confirmed in all three cell lines (Supplementary Fig. 29,30). To evaluate the intracellular cargo delivery efficiency of the peptide capsules, EGFP protein was loaded as model protein, and its internalization was quantitatively measured by FACS. VV30 and NS36 capsules also exhibited high internalization efficiency (Fig. 6b). Furthermore, the ICP capsules effectively delivered and released the hydrophilic drug Doxorubicin hydrochloride (DoXHCl). Dose-dependent viability decrease confirmed the efficient uptake and action of DoXHCl in cells (Fig. 6c). In vitro experiments demonstrated that the peptide capsules could rapidly release drugs under lower pH conditions (Supplementary Fig. 31). Confocal microscopy images illustrated the gradual release of DoXHCl from lysosomes to the nucleus after 12 hours (Fig. 6d and Supplementary Fig. 32). Additionally, the co-delivery of fluorescence-labeled BSA with DoXHCl was tested, revealing clear colocalization of DoXHCl and BSA in the lysosomes, along with released DoXHCl signal in the nucleus after 4 hours incubation (Fig. 6e and Supplementary Fig. 33). These cell experiments demonstrate the high loading and cellular delivery efficiency of the ICP capsules.

**Fig. 6.**
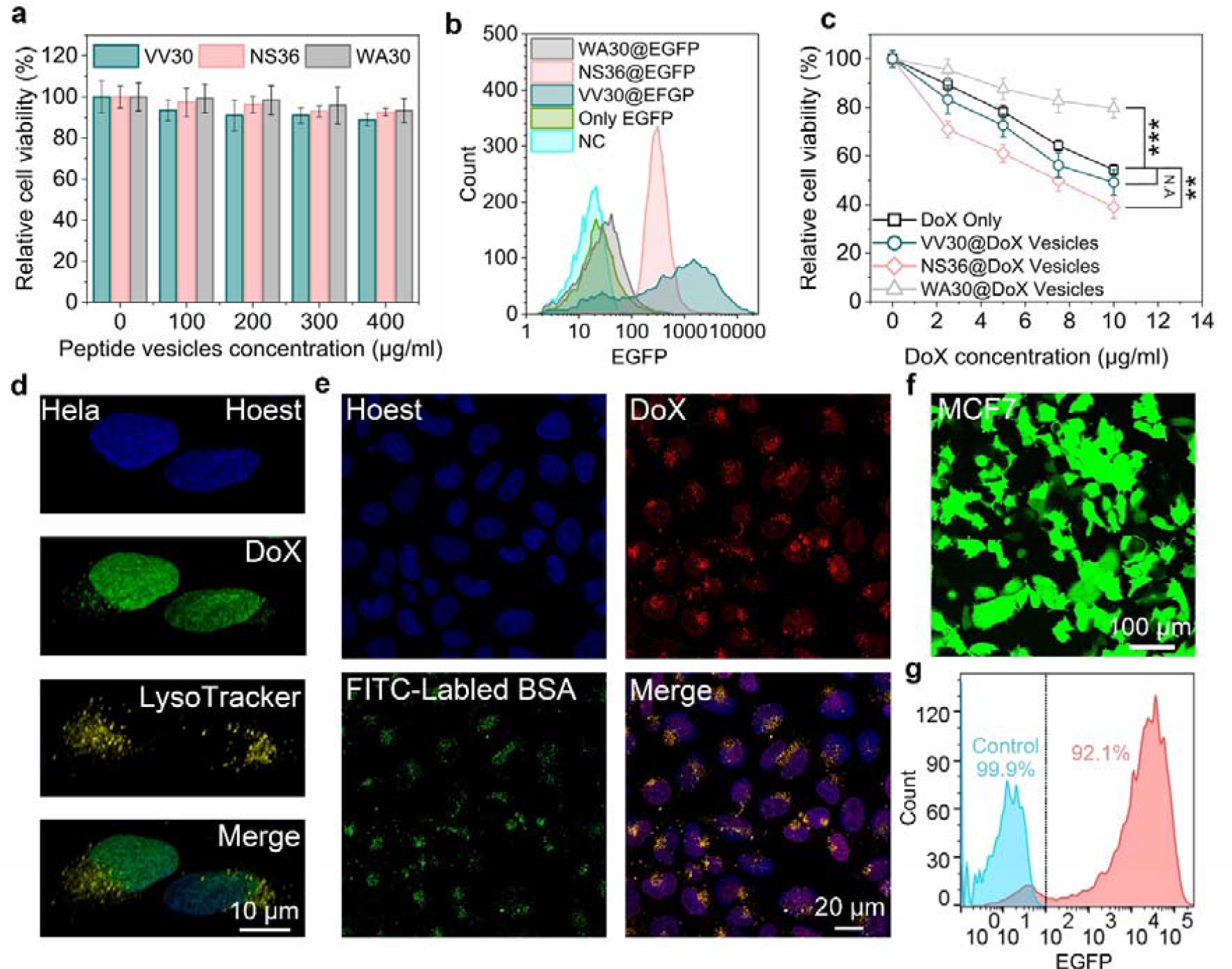
Intracellular drug, protein and mRNA delivery into cells using ICPs capsules. **a**, Cytotoxicity assessment of unloaded ICPs capsules. Data are displayed as mean□±□SD of n□=□5 independent measurements. **b**, Flow cytometry analysis of EGFP delivery mediated by ICPs capsules into Hela cells after a 2-hour incubation period. Two groups were included as controls: untreated cells (negative control, NC) and cells treated solely with EGFP without any carriers (blank). **c**, Concentration-dependent cytotoxicity of DoX and DoX-loaded ICPs capsules. Data are presented as mean□±□SD of n□=□3 independent experiments. Two-sided Student’s t-test results are shown: N.A. = 0.1960, *P□=□0.0063, ***P = 0.0007 compared to DoX alone at 10□μg/ml. **d**, 3D confocal microscopy images of Hela cells treated with DoXHCl-loaded NS36 capsules for 12 hours. This delivery result was confirmed in n□=□3 independent assays. Nuclear stain Hoechst is blue, DoX is represented in green, and lysosomal stain LysoTracker is yellow. **e**, Confocal microscopy images of Hela cells treated with FITC-labeled BSA and DoXHCl co-loaded NS36 capsules for 4 hours. This delivery result was confirmed in n□=□3 independent assays. Nuclear stain Hoechst is blue, DoXHCl is represented in red, and FITC-labeled BSA in green. **f-g**, Confocal microscopy images (f) and flow cytometry analysis (g) of MCF7 cells transfected with EGFP mRNA loaded in CC-NK3S-CC-RGD nanocapsules.

Modification in the peptide sequences further enables the delivery of nucleic acids. In the case of EGFP delivery, the VV30 capsules escaped from the lysosome to the cytoplasm by triggering the proton sponge effect through the His residues on the capsule surface^47^ (Supplementary Fig. 34). This characteristic also allows these capsules to deliver nucleic acids. We introduced two Cys residues at both the N and C terminals of the NS36 sequence for cross-linking and stabilizing the capsule structure^48^. Additionally, three Asn residues were replaced with Lys to induce the proton sponge effect while reducing the capsule size. To enhance targeted recognition, an RGD sequence was added at the C-terminal to enhance cell adsorption^49^. Based on these design principles, the CC-N3KS-CC-RGD capsules demonstrated improved delivery depth compared to NS36 capsules and successfully delivered EGFP plasmid and mRNA into MCF7, HeLa, and U2OS cells (Fig. 6f,g and Supplementary Fig. 35). For mRNA delivery, all three types of cells exhibited over 90% expression efficiency. The high cargo loading capacity and efficiency of the ICP nanocapsules render them a potentially promising platform for immunotherapy and gene therapy^50,51^.

### Conclusion and discussion

Our research provides valuable insights into how solvent gradients influence the self-assembly and encapsulation behaviors of insect cuticle peptides (ICPs). The self-assembly process operates on distinct timescales, beginning with a rapid phase where fluid-like droplets followed by a slower β-sheet formation that results in solid-like structures. Using MD simulations and FEP calculations, we further delve into the interactions between peptides and the energetics at the gradient interface. The concentration gradient, created through solvent exchange, is the key driver for nanocapsule formation. Our TFE landscape calculations uncover the fundamental mechanisms behind ICP localization and self-assembly. The nanocapsule forming peptides show unique single-well TFE profiles with a global minimum in mixing concentration between water and acetone, guiding the peptides to localize at the gradient interface. High peptide concentration facilitates β-sheet formation in the localized zone, aiding the solidification of nanocapsules. Importantly, our experiments and simulations indicate that the initial secondary structure of peptides does not decisively determine nanocapsule formation. Instead, the differing solvation free energy plays a predominant role as peptides dissolve in the constantly changing chemical environment created by the solvent mixture.

Recognizing the limitations of our study, we acknowledge several challenges that should be addressed in future research. Firstly, through systematic point mutation of the ICPs, we can fine-tune the formation process and structure of the nanocapsules, which has not been systematically examined in this study. Secondly, while our simulation approach provides valuable insights into ICP initial contact and self-assembly, it does not capture the formation process of β-sheet structures. Simulating peptide diffusion, localization, and assembly in a co-diffusional system is beyond the current capabilities of all-atom molecular dynamics in terms of time- and length-scale. Alternative methods such as coarse-grained molecular dynamics, Monte Carlo simulations, and continuum-level modeling could be explored in future endeavors to study similar systems theoretically and computationally.

Despite these limitations, our study demonstrates a straightforward solvent exchange procedure and an extensive peptide screening strategy, leading to encapsulation of diverse cargos. The nanocapsule formation mainly depends on the solvation free energy of peptides in the mixed solvent. By properly designing the energy landscape of the peptides through sequence programming, we have achieved the control of their phase and assembly behaviors. Similarly, the presence of concentration gradients in natural systems could play a pivotal role in directing the assembly and organization of proteins at biological interfaces. Moreover, we have demonstrated such nanocapsules can be customized as vehicles for targeted therapeutics and drug delivery. These revelations underscore the potential applicability of our study across a range of fields, such as nanomedicine, biomaterials, and sustainable manufacturing.

### Online content

Any methods, additional references, Nature Portfolio reporting summaries, supplementary information, acknowledgements, peer review information; details of author contributions and competing interests; and statements of data and code availability are available at **TBA**.

## Methods Materials

For solid-phase peptide synthesis, repeat peptide, resins and Fmoc-protected amino acids were purchased from GL Biochem. N,N′-diisopropylcarbodiimide was purchased from Tokyo Chemical Industry (TCI). Acetic acid, N,N-diisopropylethylamine, 1-[bis(dimethylamino)methylene]-1H-1,2,3-triazolo[4,5-b]pyridinium 3-oxide hexafluorophosphate, piperidine, tetrahydrofuran, N,N-dimethylformamide, Dichloromethane, trifluoroacetic acid, bovine serum albumin, lysozyme, rhodamine-B, were obtained from Sigma-Aldrich. LysoTracker Red DND-99, Opti-MEM, Hoechst 34580, were purchased from Thermo Fisher Scientific. Other organic solvents, including ethyl acetate, hexane and diethyl ether were purchased from Aik Moh Paints & Chemicals Pte Ltd. Dulbecco’s modified Eagle medium (DMEM), fetal bovine serum (FBS), phosphate-buffered saline buffer (PBS 10X), were purchased from Gibco. Penicillin-Streptomycin solution was purchased from Cytiva. Live cell imaging solution was purchased from Invitrogen. Trans-Blot Turbo 0.2-μm nitrocellulose transfer packs, 4–20% Criterion TGX stain-free protein gel and Clarity and Clarity Max Western ECL substrate were purchased from Bio-Rad. EGFP, and designed proteins were expressed in Escherichia coli BL21 strain and purified with Ni-NTA His Bind resin. Hela cell lines were obtained from ATCC.

### Transcriptomic analysis of larvae cuticle and CP annotation/identification

Laboratory colonies of *O. furnacalis* were originally obtained from the Institute of Plant Protection, Chinese Academy of Agricultural Sciences. Total RNAs were extracted from the head capsule and integument of the fifth instar day 1 larvae, respectively, when the head capsules were not heavily tanned. Then, the total RNAs were used for mRNA preparation and cDNA library construction.

### Peptide synthesis and purification

The original sequence peptides were customized from GL Biochem. All those peptides were purified using preparative high-performance liquid chromatography. The purified peptides were tested by positive mode electrospray ionization mass spectrometry (ESI-MS) and semi analytical and preparative HPLC. For the other mutations, the classical Merrifield solid-phase peptide synthesis (SPPS) technique was used.

### Preparation and characterization of peptide-based nanocapsules

The organic phase was added to peptide solutions at various volume ratios with vigorous stirring to generate capsules. The average hydrodynamic diameters (Dh) and Zeta potentials of peptide-based nanocapsules were measured by DLS on a Nano Particle Analyzer SZ-100 (HORIBA, Japan) at ambient temperature. Cursory morphological evaluation of the nanocapsules was performed by AFM. All the AFM images were acquired by contact (tapping) mode on an Asylum Research Cypher S with silicon nitride hard tips. Detailed morphological evaluation of the capsules was performed by TEM, those images were obtained on a Carl Zeiss Libra 120 Plus 120kV.

### Secondary structure characterization during nanocapsule formation

The secondary structure of peptides at different concertation was characterized using circular dichroism (CD) and Nuclear Magnetic Resonance (NMR) spectroscopy. ThT labeling and attenuated total reflectance Fourier transform infrared spectroscopy (ATR-FTIR) are used to monitor Protein secondary structure changes during structural changes during capsule formation.

### All-atom Molecular dynamics

The GROMACS package (version 2018.2) was utilized for all-atom MD simulations to explore the secondary structure, association, and self-assembly of ICPs in solution. Two peptides capable of forming nanocapsules (NS36 and WA30) and a precipitant-forming peptide (QH33) were investigated computationally. Temporal evolution of peptide-system interactions was analyzed using various built-in GROMACS tools, including distance, rms, rmsf, gyrate, sasa, and dssp. Specifically, the center-of-mass distance, RMSD, RMSF, radius of gyration, surfactant accessible surface area, and secondary structures were examined. Hydrogen bonds were evaluated between peptides and between peptides and the solvent using a cutoff scheme of 0.35 nm and 30 degrees.

### Transfer Free Energy Calculation

To assess the free energy changes associated with peptide nano-capsulation and precipitation, we employed the Free Energy Perturbation (FEP) method to calculate the free energy differences between the prescribed thermodynamics states in the process. Specifically, we evaluated the transfer free-energy difference (ΔΔG) by comparing the binding free energies of the peptide in various acetone concentrations from c=0 (water) to c=1 (pure acetone). Using the equation below:

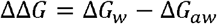

Where ΔG w and ΔG*a*w are the binding free energies of peptide in water and water-acetone mixture, respectively.

### Loading and releasing the small molecule cargoes and biomacromolecules cargo

The small molecule cargoes Rhodamine-B(RhB), FITC, Doxorubicin (DoX), Doxorubicin Hydrochloride (DoXHCl), Paclitaxel (PTX) and the macromolecule EGFP, RhB labelling BSA, EGFP-plasmid/mRNA, β-Gal and RhB-Smac were used to measure the ability of repetitive peptide capsules to transport different types of molecules. The cargo molecule and the peptide were dissolved together in deionized water and enough acetone with crosslinker solution was added dropwise. After six hours of crosslinking, the acetone solution was removed by bubbling nitrogen flow. The repeat peptide capsules were washed three times by redispersed in deionized water and collected by centrifugation to remove unreacted crosslinker and unencapsulated molecular cargo.

### Delivery of the small molecule, protein and nucleic acid cargos

For delivery protein and small molecule drugs like DoXHCl into Hela cells, the HeLa cells was treated though 950 μL Opti-MEM with 1% Penicillin-Streptomycin, then 50 μL of loaded protein capsules at 1 mg mL^-1^ was added. After 4□h of uptake, the Opti-MEM with capsules was fully removed and washed twice. The medium then was replaced by 1 mL Live Cell Imaging Solution before observed by a confocal microscope (Stellaris 5, Leica). The methylthiazolyldiphenyl-tetrazolium bromide (MTT) assay was used to evaluate the cytotoxicity of the DoX-loaded and peptide-based nanocapsules. The internalization mechanism was studied by using LysoTracker staining.

