## supplemental information for "Harnessing Gradients for Self-Assembly of Peptide-Based Nanocapsules: A Pathway to Advanced Drug Delivery Systems"

Haopeng Li[1,2], Xuliang Qian[3], Harini Mohanram[4], Xiao Han[1], Huitang Qi[2], Guijin Zou[5], Fenghou Yuan[2], Ali Miserez[1,6],Qing Yang[7], Tian Liu[2*], Huajian Gao[3,5*] and Jing Yu[1,8*]

[1]School of Materials Science and Engineering, Nanyang Technological University, 50 Nanyang Avenue, Singapore, 637553, Singapore
[2]School of Bioengineering, Dalian University of Technology, Dalian, 116024, China
[3]School of Mechanical and Aerospace Engineering, Nanyang Technological University, 50 Nanyang Avenue, Singapore, 639798, Singapore
[4]School of Biological Sciences, Division of Structural and Computational Biology, Nanyang Technological University, Singapore, 637551, Singapore
[5]Institute of High Performance Computing, A*STAR, Singapore, 138632, Singapore
[6]Biological and Biomimetic Material Laboratory (BBML), Center for Sustainable Materials (SusMat), School of Materials Science and Engineering, Nanyang Technological University, Singapore, 637553, Singapore
[7]Institute of Plant Protection, Chinese Academy of Agricultural Sciences, Beijing, 100193, China
[8]Institute for Digital Molecular Analytics and Science, Nanyang Technological University, Singapore, 637553, Singapore

**Supplementary Information**

**Methods**

**Materials.** Repeat peptide, resins and Fmoc-protected amino acids used in solid-phase peptide synthesis were purchased from GL Biochem. *N*,*N'*-diisopropylcarbodiimide was purchased from Tokyo Chemical Industry (TCI). Acetic acid, *N*,*N*-diisopropylethylamine, 1-[bis(dimethylamino)methylene]-1H-1,2,3-triazolo[4,5-b]pyridinium 3-oxide hexafluorophosphate, piperidine, tetrahydrofuran, *N*,*N*-dimethylformamide, Dichloromethane, trifluoroacetic acid, bovine serum albumin, lysozyme, rhodamine-B, were obtained from Sigma-Aldrich. LysoTracker Red DND-99, Opti-MEM, Hoechst 34580, were purchased from Thermo Fisher Scientific. Other organic solvents, including ethyl acetate, hexane and diethyl ether were purchased from Aik Moh Paints & Chemicals Pte Ltd. Dulbecco's modified Eagle medium (DMEM), fetal bovine serum (FBS), phosphate-buffered saline buffer (PBS 10X), were purchased from Gibco. Penicillin-Streptomycin solution was purchased from Cytiva. Live cell imaging solution was purchased from Invitrogen. Trans-Blot Turbo 0.2-μm nitrocellulose transfer packs, 4–20% Criterion TGX stain-free protein gel and Clarity and Clarity Max Western ECL substrate were purchased from Bio-Rad. EGFP, and designed proteins were expressed in *Escherichia coli* BL21 strain and purified with Ni-NTA His Bind resin. Hela cell lines were obtained from ATCC.

**Transcriptomic analysis of larvae cuticle and annotation of CPs.** Laboratory colonies of *O. furnacalis* were originally obtained from the Institute of Plant Protection, Chinese Academy of Agricultural Sciences. *O. furnacalis* were reared on an artificial diet at 26 to 28 °C and a relative humidity of 70% with a photoperiod of 16 h light and 8 h darkness. Total RNAs were extracted from the head capsule and integument of the fifth instar day 1 larvae, respectively, when the head capsules were not heavily tanned. Then, the total RNAs were used for mRNA preparation and cDNA library construction. This was followed by sequencing on an Illumina HiSeq 2500 platform using paired-end 150 bp reads. The raw reads were processed using Cutadapt[1] and *de novo* assembled using Trinity[2] to obtain the unigenes. The unigenes were searched against the non-redundant protein database using BLASTx with a cutoff e-value of $1 \times 10^{-5}$. The putative

CP genes were identified on the basis of the NCBI reference sequence database (https://www.ncbi.nlm.nih.gov/genbank/).

**Tandem repeated peptides searching and sequence analysis.** A mathematic program was constructed for repeated sequences that meet a specific criterion (The repeat unit contains 5 residues or more and the unit repeats at least 3 times).

**Peptide synthesis and purification.** The original sequence peptides were customized from GL Biochem. All those peptides were purified using preparative high-performance liquid chromatography (HPLC, Column: Gemini-NX 5 μm C18 110 Å, 4.6*250 mm) in a gradient of acetonitrile and water with 0.1% trifluoroacetic acid (TFA). The purified peptides were tested by positive mode electrospray ionization mass spectrometry (ESI-MS) and semi analytical and preparative HPLC. For the other mutations, the classical Merrifield solid-phase peptide synthesis (SPPS) technique was used. Wang resin was swelling in N,N-dimethylformamide (DMF) first before using. All N-terminal protected amino acid (Fmoc-AA-OH) was dissolved in DMF before coupling, than 1-[bis(dimethylamino)methylene]-1H-1,2,3-triazolo[4,5-b] pyridinium 3-oxide hexafluorophosphate (HATU, 2 equiv., 1.12mmol) and N,N-Diisopropylethylamine (DIPEA, 5 equiv.,2.80 mmol) was added into the solution. After 5 mins reaction, the mixture was added to the Wang resin in DMF at room temperature for more than 1 h coupling reaction with bubbling nitrogen flow. The resin was washed by DMF three times before using. The amino acids were coupling by Liberty Blue Automated Microwave Peptide Synthesizer from the C-terminal to N-terminal. For deprotection of the N-terminal and the side chain, 20% of piperidine and DMF mixture was added to the filtered resin for 1h at room temperature under nitrogen flow, then the resin was washed by the DMF three times to remove the piperidine. After finished all amino acids coupling, the peptide needed to be cleaved from the resin by using a cocktail containing 95% of trifluoroacetic acid (TFA), 2.5% of $H_2O$ and 2.5% of triisopropylsilane (TIPS). After being cleavage for two hours, the resin was removed by filtration, then the supernatant was evaporated to half the original volume by bubbling nitrogen flow. Impurified peptides was precipitated by adding ten times the volume of cold diethyl ether to the supernatant. The precipitation was collected by centrifugation and was fully dissolved with 10% acetic acid and 90% DI water. The HPLC (1260 prime II Infinity, Agilent Technologies) quipped with a C8 column (Zorbax 300SB-C8, Agilent Technologies) was used to purification those mutation peptides by using a gradient mixture of acetonitrile and water. The purified peptides were isolated by lyophilization (Labconco) from HPLC elutes.

**Preparation and characterization of peptide-based nanovesicles.** Peptides were dissolved in water (5 to 30 mg $mL^{-1}$ concentration with 2.5 mg $mL^{-1}$ as interval) at ambient temperature, whereas slightly excessive amount of isophorone diisocyanate (IPDI) were solved in acetone. The organic phase was poured directly into peptide solutions with ratios from 9:1 to 1:1 with vigorous stirring. After 6 h, equal volume of water was added to the mixture to quench the reaction and the acetone in the mixture was removed by gentle nitrogen flow. Then the samples were dialyzed in water for 2 days and freeze-dried into powder.

The average hydrodynamic diameters (Dh) and Zeta potentials of peptide-based nanovesicles were measured by DLS on a Nano Particle Analyzer SZ-100 (HORIBA, Japan) at ambient temperature. The freshly prepared pristine were dispersed to a concentration of 2 mg $mL^{-1}$ in water which had already adjusted pH covered 3.0 to 7.4 with 1.0 interval by 0.01 M HCl or 0.01 M NaOH.

Cursory morphological evaluation of the vesicles was performed by AFM. All the AFM images were acquired by contact (tapping) mode on an Asylum Research Cypher S with silicon nitride hard tips. All vesicles solution samples at 2 mg $mL^{-1}$ were dropped on freshly cleaved mica surface and dried under low vacuum drying oven at ambient temperature overnight. All images were fixed zero, flattened to remove background curvature, changed the stretch colour range, and cut into same size by Gwyddion 2.52, no further modify was made. Detailed morphological evaluation of the vesicles was performed by TEM, those images were obtained on a Carl Zeiss Libra 120 Plus 120kV. Vesicles samples were prepared by dropping 4 μL of dialysed sample solution with 2 mg $mL^{-1}$ of long peptide vesicles 5 mg $mL^{-1}$ of short peptide vesicles on a 300-mesh carbon-coated copper grid. The loaded copper mesh was placed in a drying oven with ambient temperature for 24 hours to let DI water evaporate. All images were taken by OLYMPUS CCD camera system under bright field with objective aperture inserted.

**Secondary structure characterization during vesicles formation.** The secondary structure of peptides at different concertation before assembly was characterised using circular dichroism. All peptides were dissolved at 0.5 to 10 mg $mL^{-1}$ concentration in DI water and loaded in a quartz cuvette with 0.2 mm path length. The data was acquired by an AVIV 420 Circular Dichroism spectrometer equipped with a temperature controller. Data were collected at 25°C over a wavelength range of 185 to 260 nm with a 1 nm wavelength steps size. Each data was averaged over three scans and smoothed via OriginLab Pro 2021.

ATR-FTIR spectra of the peptides in the self-assembling microenvironment (water was replaced by deuterium oxide) were obtained using a Bruker Vertex 70 spectrometer equipped with a PIKE Technologies MIRacle Attenuated Total Reflectance $CaF_2$ crystal reflection accessory (Bruker). The data was collected at ambient temperature over the range of 400 to 4000 $cm^{-1}$ with a resolution of 4 $cm^{-1}$. Each 10 μL sample solution was injected to the $CaF_2$ crystal surface and averaged over 64 scans. All data ware further processed by OPUS 6.5.

ThT labelling strategies were used to demonstrate structural changes during vesicle formation. ThT solution (1 mg $mL^{-1}$ in stock, further diluted by a factor of 50 to 2 μg $mL^{-1}$ as working solution) was used to dissolve peptide, then incubated by dry bath at 10 degrees. The fluorescence intensity was then tested immediately, after six hours of crosslink reaction and lyophilization then re-dissolution by DI water, then recorded the fluorescence intensity. The ThT fluorescence data was collected by a Cary Eclipse Spectrophotometer (Agilent) at excitation 450 nm and emission 485 nm. The optical microscope and fluorescence labelled vesicle images were captured by Olympus IX73 inverted microscope with a CoolLED-pE-300-Ultra-LED-Fluorescence-Light-Source accessory.

**Nuclear Magnetic Resonance (NMR) spectroscopy.** The NMR experiments were carried out in 600 MHz spectrometer equipped with cryoprobe. Two dimensional $^{1}H$-$^{1}H$ TOCSY (Total Correlation spectroscopy) and $^{1}H$-$^{1}H$ NOESY (Nuclear Overhauser Spectroscopy) measurements were made for 0.5 mM of peptides (QH33, WA30 and NS36) in water, pH 4.5 with 10% $D_2O$ for deuterium lock and DSS for signal reference with 80 ms and 200 ms mixing times, respectively. Uniform $^{13}C$ and $^{15}N$ labelled NS36 peptide was prepared as discussed above and the backbone NMR spectra viz HNCA, HNCACB and CACBCONH along with $^{1}H$-

$^1H$-$^{15}N$ NOESY-HSQC spectra were also acquired for further analysis.

The molecular level conformational landscape information was obtained by analyzing 2D $^1H$-$^1H$ TOCSY and $^1H$-$^1H$ NOESY experiments of QH33, WA30 and NS36. Of the three peptides, the number of NOE connectivities were observed in order of WA30>NS36>QH33 proposing structural propensities in WA30 and NS36.

For WA30 peptide, all of the residues can be unambiguously assigned. $^1H^a$ chemical shift deviations of non-proline residues exhibited negative trend suggesting alpha helical conformation. A closer observation of $^1H$-$^1H$ NOESY spectrum revealed a number of *i,i+3* and *i,i+4* NOEs spanning over residues S16-A28, diagnostic of helical conformation. The three-dimensional structure of WA30 was then calculated using all of these medium range NOEs that culminated into partially folded a-helix in the C-terminus. The helical fold in the N-terminus was found to be interrupted by the presence of P13 and P15 residues. This flexibility in turn brought the side chain of the residues W9 and W16 closer to be involved in π-π interaction and the side chains of W25 were found to be exposed. Thus, it can be speculated that the π-π interaction of W9-W16 stabilizing monomeric conformation whereas the exposed W25 can interact with other monomers bringing about the formation of nanoparticles.

For NS36 peptide, the presence of two Pro residues sequentially affected the sequential walk of TOCSY and NOESY spectra. But since we were successful in expressing and purifying uniform isotope ($^{13}C$, $^{15}N$) labelled peptide, 3D backbone experiments (HNCA, HNCACB and CACBCONH) assisted the complete assignment of the peptide. The 2D $^1H$-$^{15}N$ HSQC spectrum was well dispersed over a range of 7.5-9.0 ppm. The positive and negative chemical shift deviations of $^{13}Ca$ and $^1Ha$ indicated helical propensity for NS36 for all non-proline residues. The diagnostic *i,i+3* and *i,i+4* peaks were also found to be present but sparsely located when compared to WA30. The three-dimensional structure calculated with all of the assignments resulted in flower like loop structure. A closer observation revealed the presence of π-π side chain interaction between Y11 and Y18 residues with flexibility support from P13 and P14 residues as highlighted with different shades of green. On the other hand, the side chains of rest of the Tyr residues were exposed which might interact with other Tyr residues when they form nanoparticles.

For QH33 peptide, all of the residues can be assigned unambiguously, and the positive chemical shift deviations indicated an extended conformation. The 2D $^1H$-$^1H$ NOESY spectrum of QH33 also displayed very limited NOE connectivities mainly between backbone. Hence the three-dimensional structure of QH33 displayed an extended conformation with all of the sidechains of His residues exposed.

Overall, the structural studies of monomeric conformations of WA30, NS36 and QH33 peptide explained the molecular mechanism behind sequence specific nanoparticle formation. In the case of WA30 and NS36 peptides, the helical landscape was mostly interrupted by the presence of consecutive or closely available Pro residues that in turn provided the flexibility for the aromatic residues (Trp in WA30 and Tyr in NS36) to engage in π-π side chain interactions. On the other hand, because of the absence of intervening Pro residues in QH33, there is no structural stability of monomers that can bring about the formation of higher order structures and thus nanoparticle formation.

**All-atom Molecular dynamics.** The GROMACS package (version 2018.2) was utilized for all-atom MD simulations to explore the secondary structure, association, and self-assembly of ICPs in solution. Two peptides capable of forming nano-capsules (NS36 and WA30) and a precipitant-forming peptide (QH33) were investigated computationally. The initial peptide configuration were obtained through liquid-phase NMR experiments. Following the protocol from our previous study[3], the amber99ff99SB force field was employed for the MD simulations, including the assignment of partial charges and force field parameters[4]. The TIP3P water model was employed[5], and the general Amber force field (GAFF) parameters for acetone were adopted[6]. The same acetone force field was reported in literature for the study of biphasic mixing[7]. In the QH33 simulation, counterions ($Na^+$) were introduced to neutralize the system. The LINCS algorithm was used to constrain hydrogen atoms[8]. To simulate a water-acetone mixed solution, the simulation box, with dimensions of 10 × 10 × 20 nm, was initially filled with water molecules in half of the simulation cell and pre-relaxed acetone molecules in the other half. The solvent system was relaxed through a short steepest-descent energy minimization, followed by a 300 ps NVT equilibrium simulation with a force constant of 1000 kJ/mol/nm applied to the heavy (non-hydrogen) atoms (oxygen for water and carbon/oxygen for acetone). Subsequently, a 300 ps NVT equilibrium simulation was conducted without restraints. The equilibrated structures were then subjected to a 50 ns molecular dynamics simulation to mix water and acetone, while monitoring the water/acetone concentration along the reaction coordinate. An in-house Tcl script was employed to extract cubic boxes of water-acetone mixtures at the desired target concentrations, along with a pre-relaxed pure water box of the same size. The resulting six sub-systems (vA/vW = 0, 1, 3, 5, 7, 9) were duplicated in the x, y, and z directions to expand the simulation boxes to a size of approximately 10 nm. These systems were then subjected to a 100 ns relaxation in the NPT ensemble to reach equilibrium. The relaxed systems were subsequently used to solvate the peptides using the GROMACS solvate command. The resulting simulation systems were sufficiently large to prevent peptide molecules from interacting with their periodic images. A time step of 2 fs was employed, and the systems were simulated in the NPT ensemble at 1 atm and 300 K. Periodic boundary conditions were imposed in all three directions. Temperature and pressure were maintained using the Nose-Hoover and Parrinello-Rahman algorithms, respectively.[9,10]

To investigate the self-assembly of peptides in different chemical environments, the system was duplicated in the x-direction to create dual-peptide systems and duplicated in the xy-directions to generate quadra-peptide systems. These multi-peptide simulations underwent the same equilibrium procedure as single peptides, with the only difference being longer relaxation times (ranging from 100 ns to 250 ns) to ensure sufficient interactions and preliminary peptide self-assembly. Temporal evolution of peptide-system interactions was analysed using various built-in GROMACS tools, including distance, rms, rmsf, gyrate, sasa, and dssp. Specifically, the root-mean-square deviation (RMSD), root-mean-square fluctuation (RMSF), radius of gyration (Rg), surfactant accessible surface area (SASA), and secondary structures were examined. Hydrogen bonds were evaluated between peptides and between peptides and the solvent using a cutoff scheme of 0.35 nm and 30 degrees.

**Transfer Free Energy Calculation.** To assess the free energy changes associated with peptide nano-capsulation and precipitation, we employed the Free Energy Perturbation (FEP) method to calculate the free energy differences between the prescribed thermodynamics states in the process. Specifically, we evaluated the transfer free-energy difference ($\Delta\Delta G$) by comparing the binding free energies of the peptide in various acetone concentrations from c=0 (water) to c=1 (pure acetone). The transfer free-energy difference can be calculated as

$$\Delta\Delta G = \Delta G_w - \Delta G_{aw}$$

Where $\Delta G_w$ and $\Delta G_{aw}$ are the binding free energies of peptide in water and water-acetone mixture, respectively. To initiate the FEP calculations, we utilized starting configurations derived from single peptide Molecular Dynamics (MD) simulations, ensuring a reliable foundation for our FEP analysis. A time step of 1 fs and a total of 25 intermediate states were used, aligned with previous studies[11]. Each intermediate state was sampled for a duration of 4 ns. The Multistate Bennett Acceptance Ratio (MBAR) method was employed to compute the change in free energy associated with the annihilation process, using the open-source Alchemical Analysis script[12].

**Loading and releasing the small molecule cargoes and protein cargo.** The small molecule cargoes Rhodamine-B (RhB), FITC, Doxorubicin (DoX), Doxorubicin Hydrochloride (DoXHCl), Paclitaxel (PTX) and the macromolecule EGFP, RhB labelling BSA, EGFP-plasmid/mRNA, β-Gal and RhB-Smac were used to measure the ability of repetitive peptide vesicles to transport different types of molecules. The cargo molecule and the peptide were dissolved together in deionized water and a sufficient amount of acetone with crosslinker solution was added dropwise. After six hours of crosslinking, the acetone solution was removed by bubbling nitrogen flow. The repeat peptide vesicles were washed three times by redispersed in deionized water and collected by centrifugation to remove unreacted crosslinker and unencapsulated molecular cargo. Molecular cargo loading was determined by the UV-Vis-NIR Cary 5000 (Agilent). The encapsulation efficiency was calculated as

$$\frac{m_t - m_u}{m_t} \times 100\%$$

Where $m_t$ and $m_u$ represent the mass of the small molecule cargoes used and the mass of the unloading cargoes, respectively.

The RhB labelling BSA release was conducted by a dialysis bag (10kDa). The freeze-dried vesicles (1mg) were first redispersion into PBS (10 mL, pH 7.4, ionic strength 0.15M) and dialysed against 200 mL PBS (pH 7.4, ionic strength 0.15M). The protease (2mg, bovine

pancreas) was added to the vesicles suspension after dialysing 3h. Each 20 minutes, a 200 μL sample was took from the outer dialysis PBS to measure the release of RhB labelling BSA by using a microplate reader using 560nm / 610nm for the excitation/emission wavelengths.
The small molecule cargoes was also conducted by a dialysis bag (10kDa). The freeze-dried vesicles (1mg) were first redispersion into PBS (10 mL, pH 7.4, ionic strength 0.15M) and dialysed against 200 mL PBS (pH 7.4, ionic strength 0.15M). Each 1 hour, a 200 μL sample was took from the outer dialysis PBS to measure the release of small molecules by using a microplate reader using 560nm / 610nm for the excitation/emission wavelengths and UV–Vis spectra absorbance at 488nm, respectively.

**Cytotoxicity study.** The methylthiazolyldiphenyl-tetrazolium bromide (MTT) assay was used to evaluate the cytotoxicity of the DoX-loaded and peptide-based vesicles. $1 \times 10^4$ HeLa cells in 100 μL of cell culture medium (High Glucose DMEM, 5% FBS, 1% Penicillin-Streptomycin) were transferred into a 96-well plates and incubated for 24 h. The pH 7.20 1X PBS was used to wash each well twice and then replaced with 100 μL of Opti-MEM containing various concentrations of peptide-based vesicles and the DoX-loaded vesicles (various concentration of DoX, 0.5 mg $mL^{-1}$ peptide-based vesicles). After 4 h of uptake, the Opti-MEM was fully removed and the cells were washed by pH 7.20 PBS twice before changing back to 100 μL of cell culture medium (High Glucose DMEM, 5% FBS, 1% Penicillin-Streptomycin). The 96-well plates containing HeLa cells was incubated for another 24 h before 10 μL of 5 mg $mL^{-1}$ MTT dissolved in pH 7.20 PBS was added to each well. After another 4 h of incubation, all medium was removed, and the cells were washed by pH 7.20 PBS twice. Then, 100 μL of DMSO was added to each well and mixed evenly through the pipette. The absorbance at 570 nm was measured by a microplate reader (Infinite M200 Pro, Tecan). The relative cell viability was calculated as

$$\frac{\boldsymbol{A_s - A_b}}{\boldsymbol{A_{nc} - A_b}} \times \mathbf{100\%}$$

where $A_s$, $A_b$ and $A_{nc}$ represent the absorbance of the sample cells, blank control, and negative control cells, respectively.

**Delivery of the small molecule cargoes and protein cargos.** For delivery protein and small molecule drugs like DoXHCl into Hela cells, the total amount of $1\times 10^5$ HeLa cells was uniformly dispersed by Trypsin in 1 mL cell culture medium (High Glucose DMEM, 5% FBS, 1% Penicillin-Streptomycin), and then subculture into a 350 $mm^2$ glass bottom petri dish. After incubation for 24 h, the medium was changed into 950 μL Opti-MEM with 1% Penicillin-Streptomycin, then 50 μL of loaded protein vesicles at 1 mg $mL^{-1}$ was added. After 4 h of uptake, the Opti-MEM with vesicles was fully removed and the cells were washed by pH 7.20 PBS twice before adding back to culture medium (High Glucose DMEM, 5% FBS, 1% Penicillin-Streptomycin). Those cells were incubated for another 12 hours and washed again by pH 7.20 PBS, the medium then was replaced by 1 mL Live Cell Imaging Solution (invitrogen). The vesicles uptake was evaluated by FACS (LSR Fortessa X20, BD Biosciences) and observed by a confocal microscope (Stellaris 5, Leica).

The internalization mechanism was studied by using LysoTracker staining. After cells uptake of FITC-labeled BSA carrying vesicles, the cells were washed by pH 7.20 PBS twice before adding back to 1 mL culture medium (High Glucose DMEM, 5% FBS, 1% Penicillin-Streptomycin). After 2 hours culture, the medium was replaced by 1 mL Opti-MEM containing 50 nM LysoTracker Red DND-99. The cells were further stained in the medium for 1 hour. The staining treated HeLa cells were washed by pH 7.20 PBS twice and and fixed with 5% formaldehyde solution. The medium then was replaced by 1 mL Live Cell Imaging Solution (invitrogen) and observed by a confocal microscope (Stellaris 5, Leica) though super resolution model.

**Delivery of plasmid and mRNA.** For delivery plasmid and mRNA into different cells, the total amount of $1\times 10^5$ HeLa, MCF7 and U2OS cells was uniformly dispersed by Trypsin in 1 mL cell culture medium (High Glucose DMEM, 5% FBS, 1% Penicillin-Streptomycin), and then subculture into a 350 $mm^2$ glass bottom petri dish. After incubation for 24 h, the medium was changed into 950 μL Opti-MEM with 1% Penicillin-Streptomycin, then 50 μL of loaded plasmid or mRNA vesicles at 1 mg $mL^{-1}$ was added. After 4 h of uptake, the Opti-MEM with vesicles was fully removed and the cells were washed by pH 7.20 PBS twice before adding back to culture medium (High Glucose DMEM, 5% FBS, 1% Penicillin-Streptomycin). Those cells were incubated for another 12 hours and washed again by pH 7.20 PBS, the medium then was replaced by 1 mL Live Cell Imaging Solution (invitrogen). The vesicles uptake was evaluated by FACS (LSR Fortessa X20, BD Biosciences) and observed by a confocal microscope (Stellaris 5, Leica).

**Supplementary Table 1.** Amino acid content of the *Ostrinia furnacalis* (Asian corn borer) head shells

| | Ala (A) | Asn (N) | Cys (C) | Gln (Q) | Glu (E) | Gly (G) | His (H) | Ile (I) | Leu (L) |
|---|---|---|---|---|---|---|---|---|---|
| O4L_3_Head | 13.03% | 2.40% | 0.62% | 5.08% | 5.32% | 11.02% | 6.29% | 3.93% | 3.94% |
| O5L_1_Head | 22.00% | 2.51% | 0.52% | 4.39% | 3.41% | 7.17% | 5.18% | 2.89% | 5.19% |
| Eukaryotic | 7.40% | 4.50% | 1.70% | 4.10% | 6.40% | 6.10% | 2.40% | 5.30% | 9.20% |
| | Lys (K) | Met (M) | Phe (F) | Pro (P) | Ser (S) | Thr (T) | Trp (W) | Tyr (Y) | Val (V) |
| O4L_3_Head | 4.43% | 0.70% | 2.52% | 8.06% | 7.66% | 3.57% | 0.28% | 6.24% | 8.97% |
| O5L_1_Head | 2.76% | 0.69% | 1.93% | 8.35% | 7.53% | 3.15% | 0.67% | 5.78% | 10.20% |
| Eukaryotic | 5.70% | 2.20% | 3.90% | 5.40% | 8.60% | 5.70% | 1.20% | 3% | 6.10% |

**Supplementary Table 2.** Sequence information of insect cuticle peptides obtained through specific screening threshold

| Name | Sequences | Repeat unit | GRAVY |
|---|---|---|---|
| NS36 | NNYYVPPS<br>NNYVPPS<br>NNYVPPS<br>NNYVPPS<br>NNYVPPS | NNYVPPS | -1.161 |
| WA30 | WNAAPA NQ<br>WNAAPA PS<br>WNAAPA NH<br>WNAAPA | WNAAPA | -0.617 |
| VV30 | VHSAPV VHSAPV<br>VHSAPV VHSAPV<br>VHSAPV | VHSAPV | 0.767 |
| SF32 | SSGGGGGF<br>SSGGGGGF<br>SSGGGGGF<br>SSGGGGGF | SSGGGGGF | -0.100 |
| GL33 | GGGHSSGGLSL<br>GGGHSSGGLSL<br>GGGHSSGGLSL | GGGHSSGGLSL | 0.000 |
| AP36 | AVSYSAP<br>AVSYSAP A<br>AVSYSAP<br>AVSYSAP<br>AVSYSAP | AVSYSAP | 0.508 |
| GH42 | GHEGHDH<br>GPEGHDH<br>GHEGHDH<br>GPEGHDH<br>GHEGHDH<br>GPEGHDH | GHEGHDH<br>GPEGHDH | -2.371 |
| YK34 | YSEPAK F<br>YSEPAK I YSEPAK<br>I YSEPAK V<br>YSEPAK | YSEPAK | -0.897 |
| QH33 | QSHDGH EA<br>QSHDGH EA<br>QSHDGH QV<br>QSHDGH QE<br>QSHDGH | QSHDGH | -2.176 |

**Supplementary Table 3.** The assembly phenomena of different solvents on insect cuticle peptide phase separation

| Name | Dimethyl sulfoxide | Dimethyl formamide | Acetone | i-propanol | Tetrahydrofuran | Ethyl ether |
|---|---|---|---|---|---|---|
| WA30 | N.A | N.A. | LLPS > vesicles | N.A | LLPS > vesicles | Gel-like precipitate |
| NS36 | N.A | N.A. | LLPS > vesicles | N.A | LLPS > vesicles | Gel-like precipitate |
| VV30 | N.A | N.A. | LLPS > vesicles | N.A | LLPS > vesicles | precipitate |
| QH33 | N.A | particles | particles | N.A | particles | precipitate |
| N.A. indicates that no assembly occurs under this condition. | | | | | | |

**Supplementary Table 4**. The secondary structure stability monitoring of ICP capsules by ThT staining

| | 1 day | 7 days | 1 month | 3 months | 6 months |
|---|---|---|---|---|---|
| WA30 | 448 ± 9.31 | 492 ± 11.43 | 486 ± 15.97 | 477 ± 23.41 | 485 ± 26.57 |
| NS36 | 462 ± 7.63 | 456 ± 9.32 | 448 ± 14.15 | 432 ± 18.93 | 428 ± 19.37 |
| VV30 | 782 ± 28.12 | 813 ± 34.42 | 857 ± 38.63 | 861 ± 35.47 | 851 ± 32.08 |

**Supplementary Table 5.** The diffusion constant of water, acetone, and the system, determined from MD simulations of water-acetone diffusion

| Molecule Name | Diffusion Constant (1e-5 $cm^2/s$) |
|---|---|
| Water | 1.401 ± 0.022 |
| Acetone | 1.071 ± 0.012 |
| System | 1.258 ± 0.007 |

**Supplementary Table 6.** Simulation-derived timescales for peptide assembly initiation based on initial contact events

| $C_{acetone}$ | QH33 | | WA30 | | NS36 | |
|---|---|---|---|---|---|---|
| | Duo MD (ns) | Quad MD (ns) | Duo MD (ns) | Quad MD (ns) | Duo MD (ns) | Quad MD (ns) |
| 0 | 9 | 5 | 16 | 6 | 9 | 7 |
| 0.20 | 41 | 17 | 9 | 15 | 32 | 18 |
| 0.42 | 69 | 7 | 16*2, 120 | 30 | 54 | 20 |
| 0.55 | 36 | 46 | 44 | 47 | 61 | 14 |
| 0.63 | 9 | 42 | 40 | 71 | 32 | 44 |
| 0.69 | 148*1 | 5 | 23 | 14*3, 36 | 54 | 13 |

Duo MD indicates the dual-peptide MD systems;
Quad MD indicates the quadra-peptide MD systems;
*1 indicates that simultated duo-QH33 separated at 208 ns.
*2 indicates that simultated duo-WA30 separated at 39 ns.
*3 indicates that simultated quadra-WA30 separated at 20 ns.

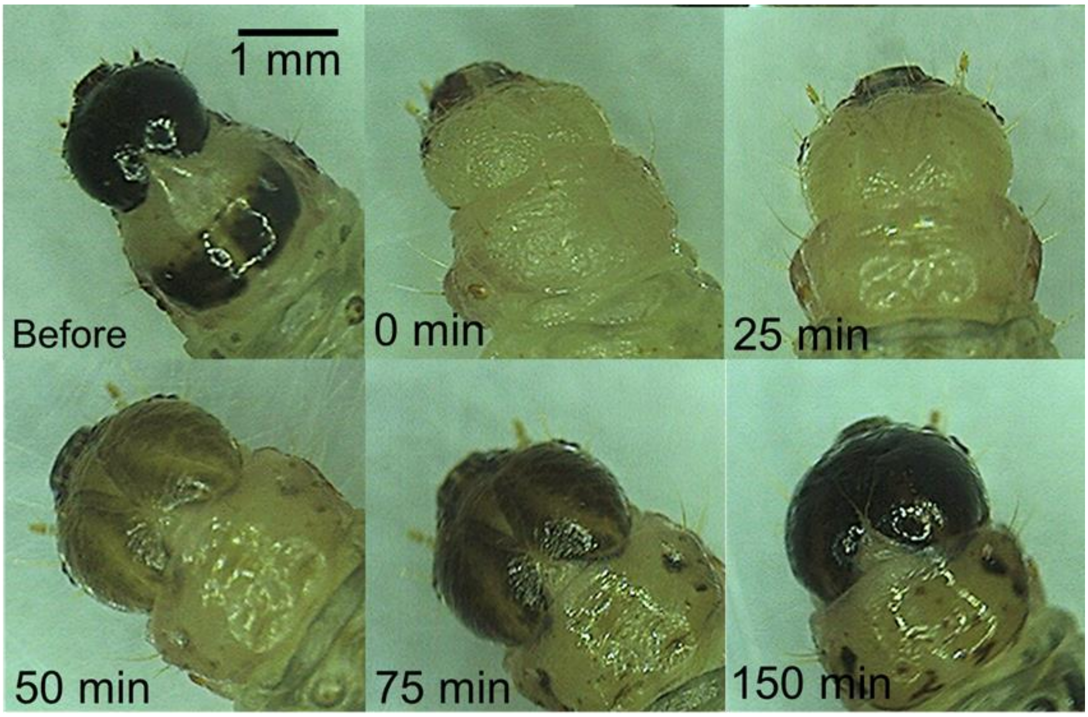


**Supplementary figure 1.** The fourth molting and capsule hardening process of the *Ostrinia furnacalis* (Asian corn borer).

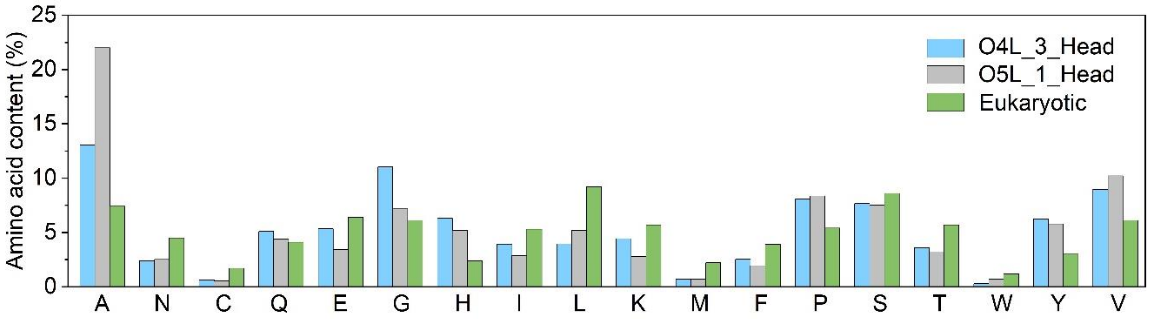


**Supplementary figure 2.** Amino acid content of the *Ostrinia furnacalis* (Asian corn borer) head shells.

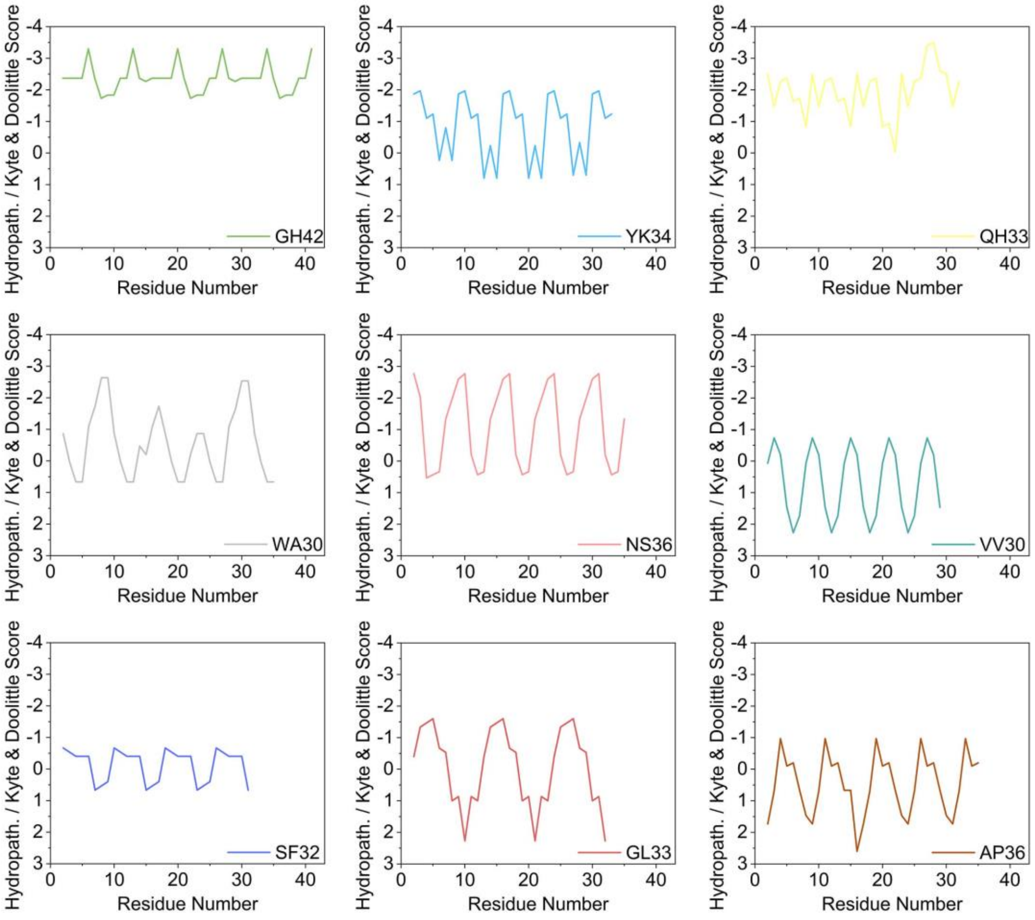


**Supplementary figure 3.** GRAVY plot of insect cuticle peptides through ProtScale by using Kyte & Doolittle scale.

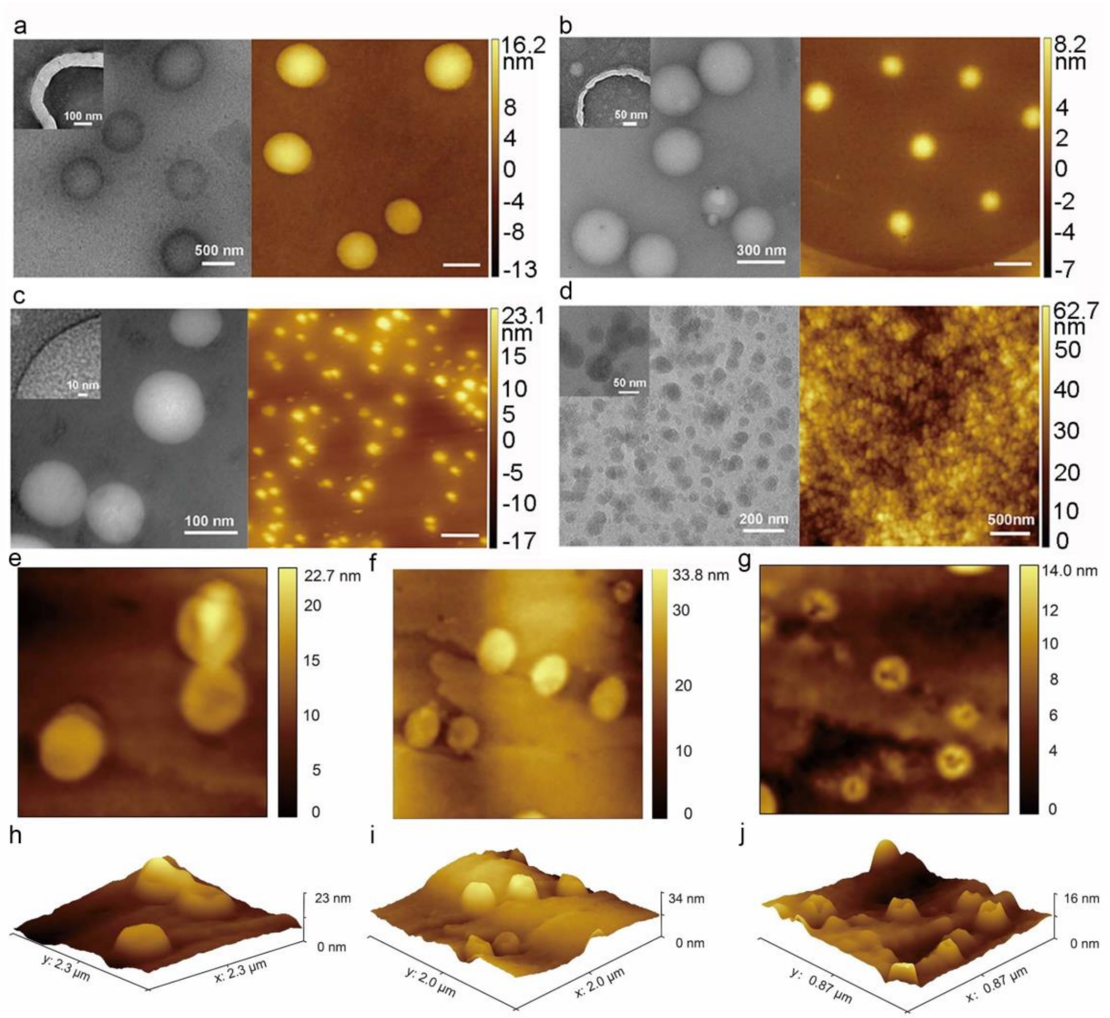


**Supplementary figure 4.** Representative AFM and 3D reconstruction images of WA30 (a,e and h), NS36 (b,f and i) and VV30 (c,g and j) cross-linked peptide vesicles and QH33 (d) cross-linked peptide particles, all scale bar have been indicated in the images, respectively.

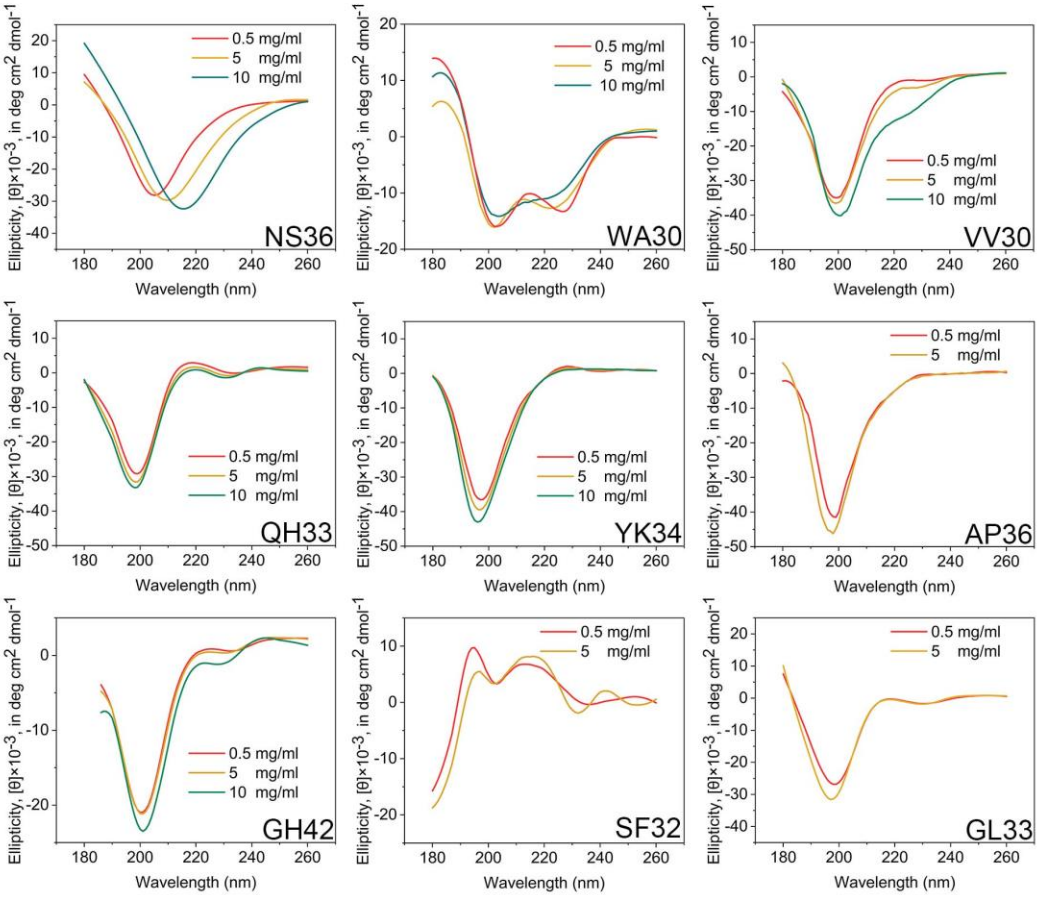


**Supplementary figure 5.** CD spectra of ICPs in aqueous solutions under different concertation.

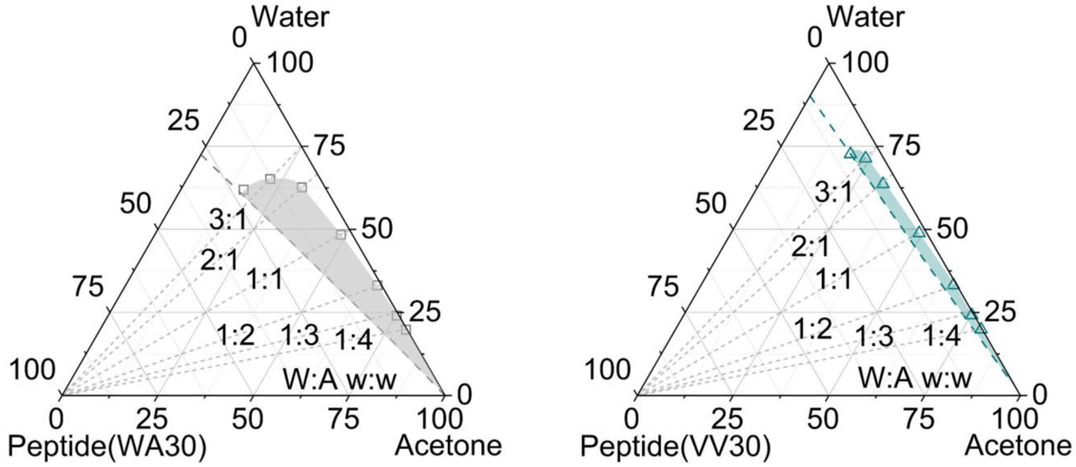


**Supplementary figure 6.** LLPS Phase diagram of WA30(a), VV30(b) in Water/Acetone Systems.

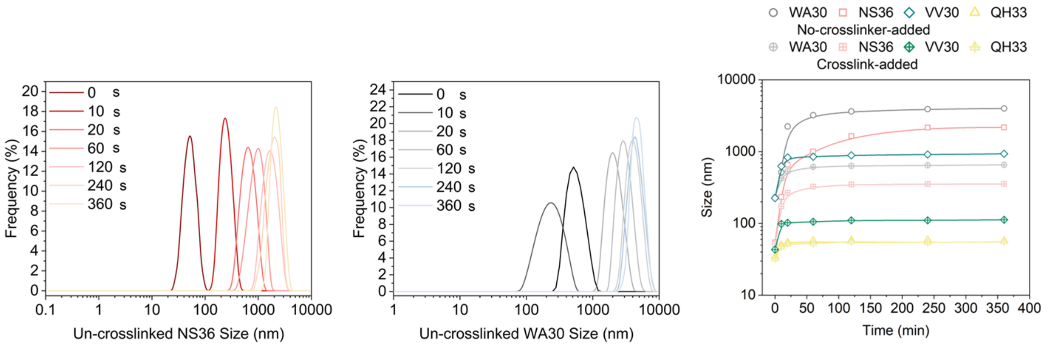


**Supplementary figure 7.** Size distributions of NS36(a), WA30(b) LLPS droplets with generation time; (c) The influence of adding crosslinking agents (IPDI) on the size of ICPs capsules.

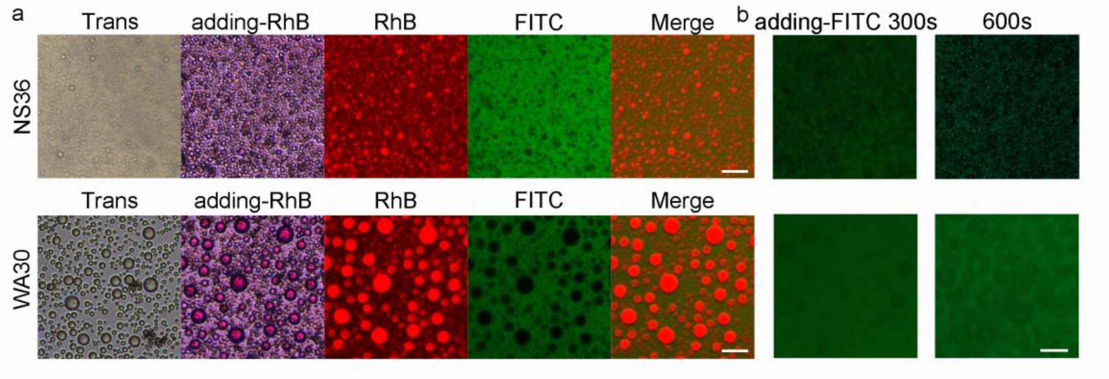


**Supplementary figure 8. a,** Fluorescence distribution micrograph of freshly prepared NS36(top) and WA30(bottom) coacervate droplets infiltrated with FITC (2 mg mL$^{-1}$ in 9 μL acetone) and Rhodamine-B (18 mg mL$^{-1}$ in 1 μL water), final mixed solution under 9:1 vol ratio of acetone to water and final peptide concertation at 4 mg mL$^{-1}$. **b**, Fluorescence microscope image of FITC distribution in the NS36 (top) and WA30 (bottom) vesicles systems changing with time. Scale bar is 10 μm.

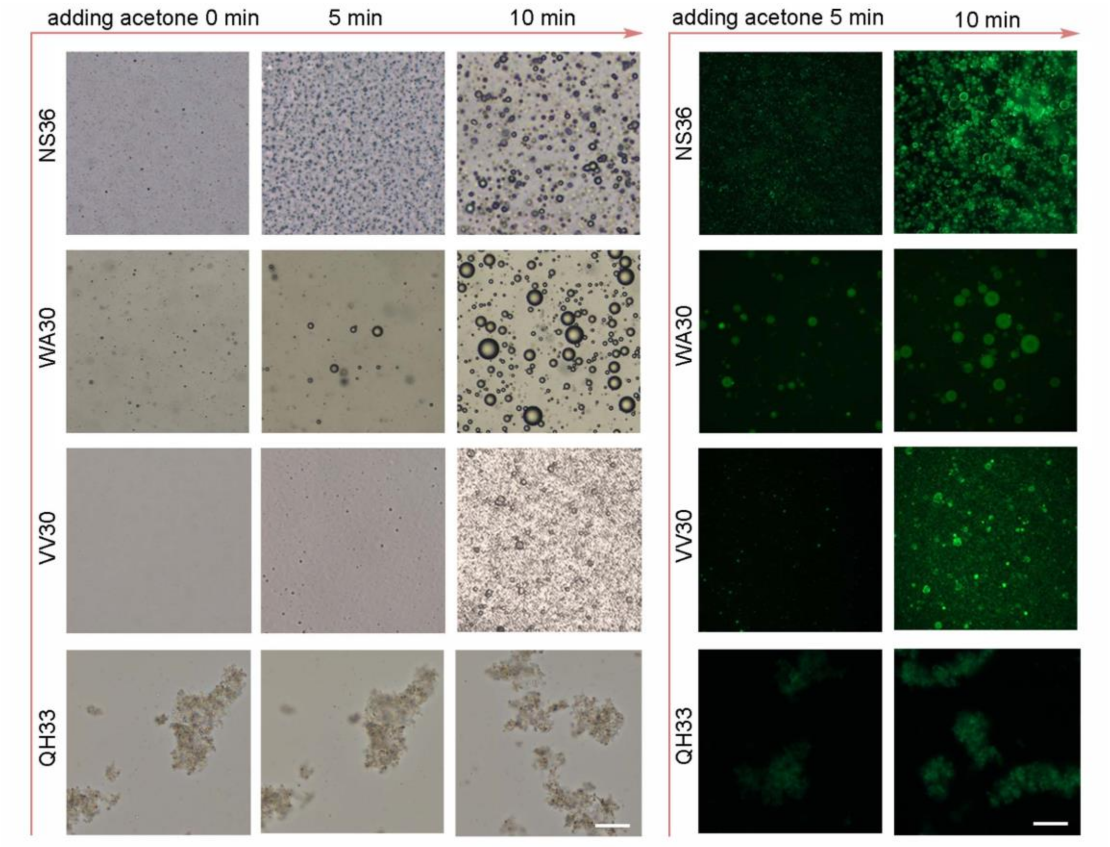


**Supplementary figure 9.** Optical (left) and fluorescence (right) micrograph of peptide LLPS droplets and vesicles (2 mg $mL^{-1}$) at 9:1 vol ratio of acetone to water within and after LLPS formation (representative images of three independent experiments). Scale bar is 5 μm.

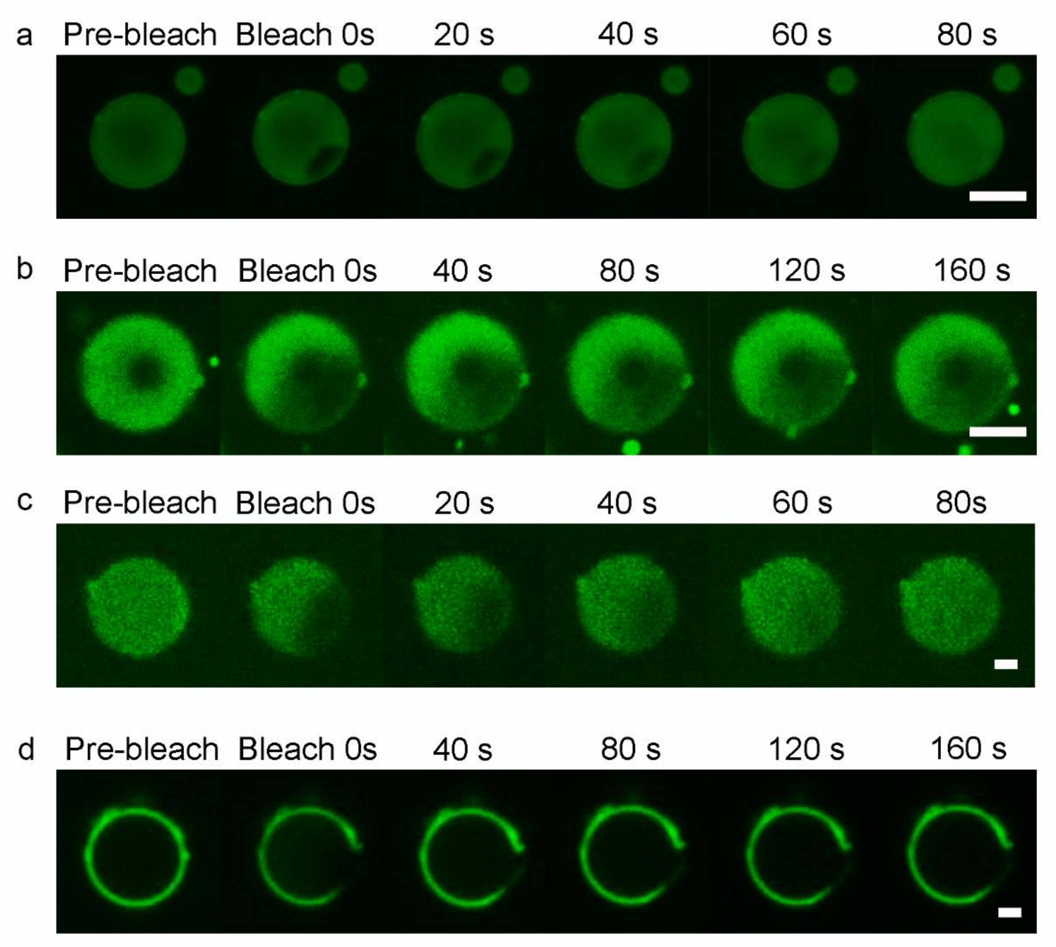


**Supplementary figure 10.** Representative confocal images during FRAP of freshly prepared coacervate droplet and vesicles of WA30 (a and b) and NS36 (c and d) under 9:1 vol ratio of acetone to water. Scale bar for a and b is 5 µm, for c and d is 1 µm.

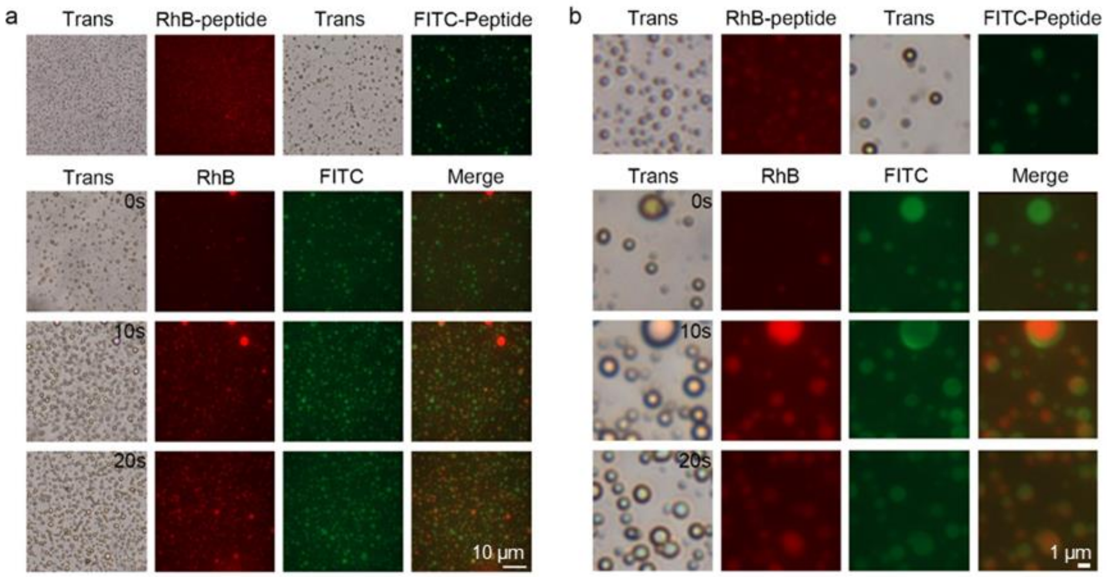


**Supplementary figure 11.** The LLPS droplets formed by FITC and RhB labeled NS36 fuse with each other after forming phase separation. (a) Image under 60x objective lens; (b) Partial zoom in image.

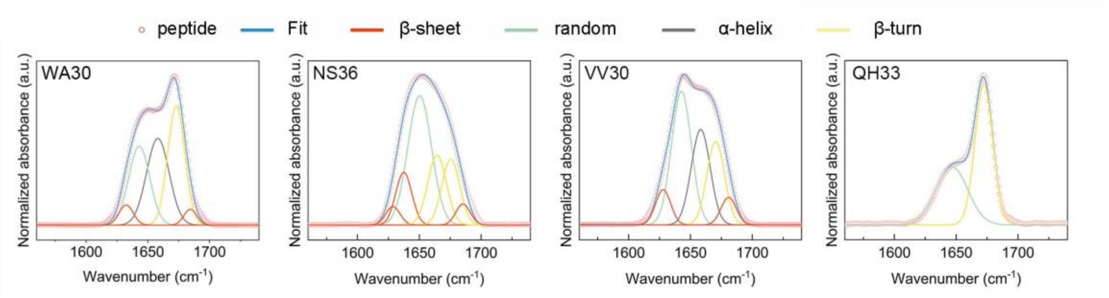


**Supplementary figure 12.** FTIR deconvolution analysis of the amide I of WA30, NS36, VV30 and QH33 in aqueous solution under 10 mg $mL^{-1}$, data are means ± SD from n = 3 independent assay.

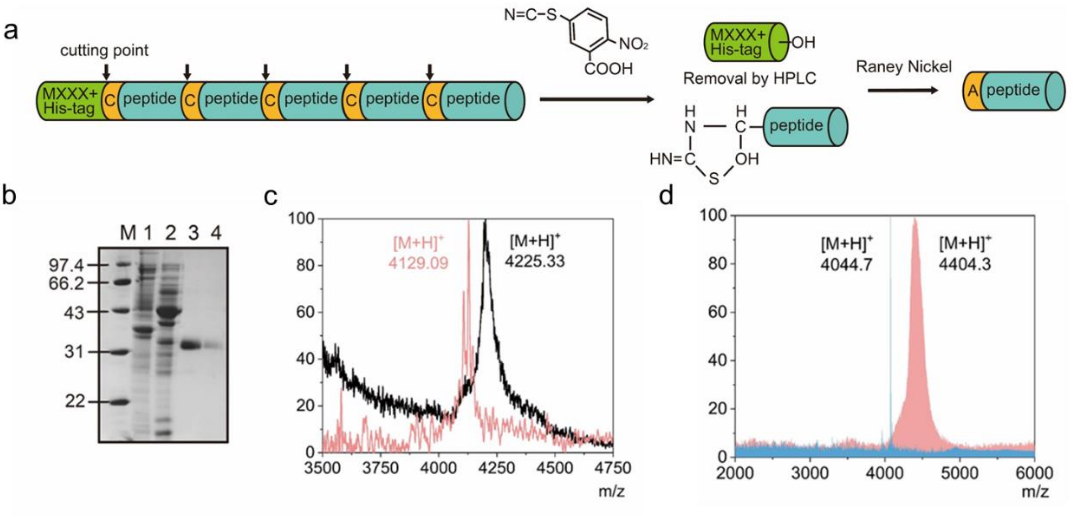


**Supplementary figure 13.** (a)Expression of isotopically substituted NS36 by the E. coli; (b) target proteins constructed by five NS36 tandem purified by His-tag, channel 1 and 2: total protein; channel 3 and 4: target protein; (c) MALD-TOF spectra of NS36 with supernumerary ring structure(black) and an additional alanine(pink); (d) MALD-TOF spectra of NS36(pink) and isotopically substituted NS36(blue).

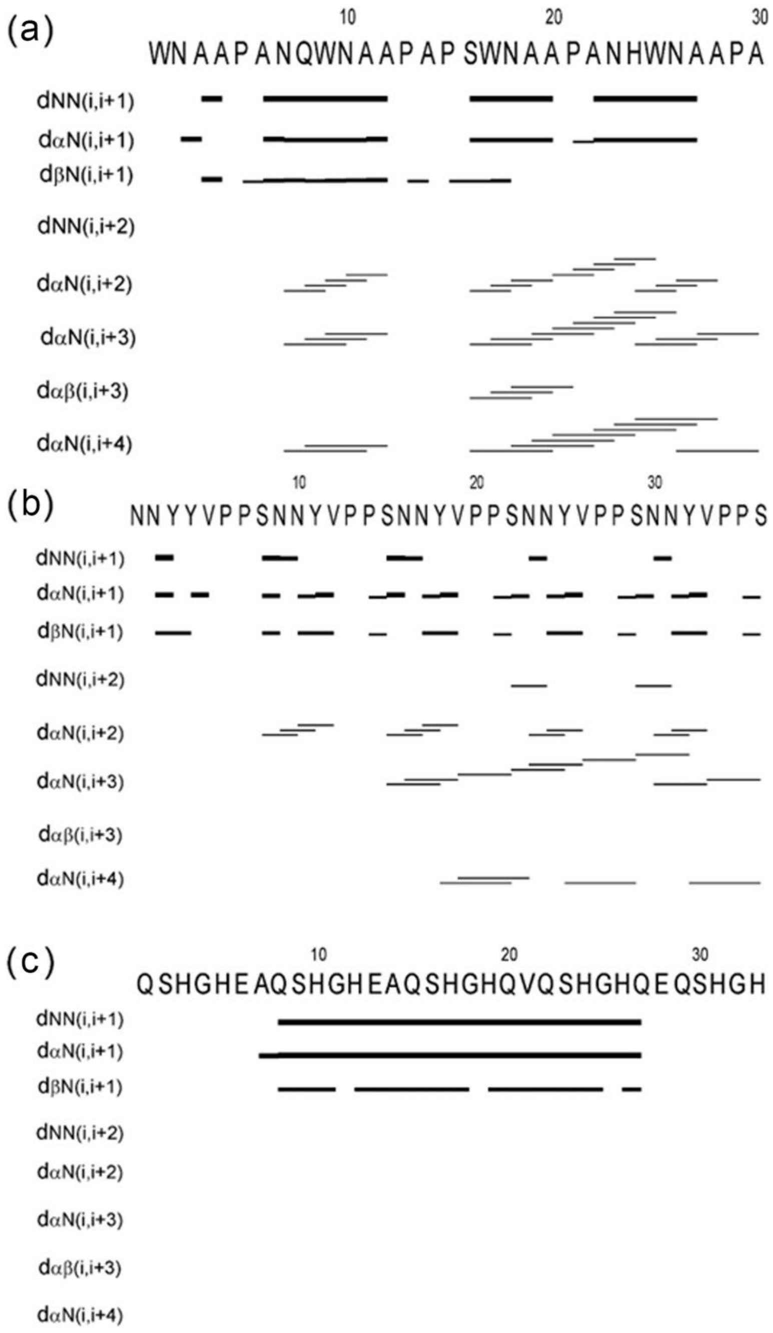


**Supplementary figure 14.** Bar diagram representations of NOE connectivities of (a) WA30, (b) NS36 and (c) QH33.

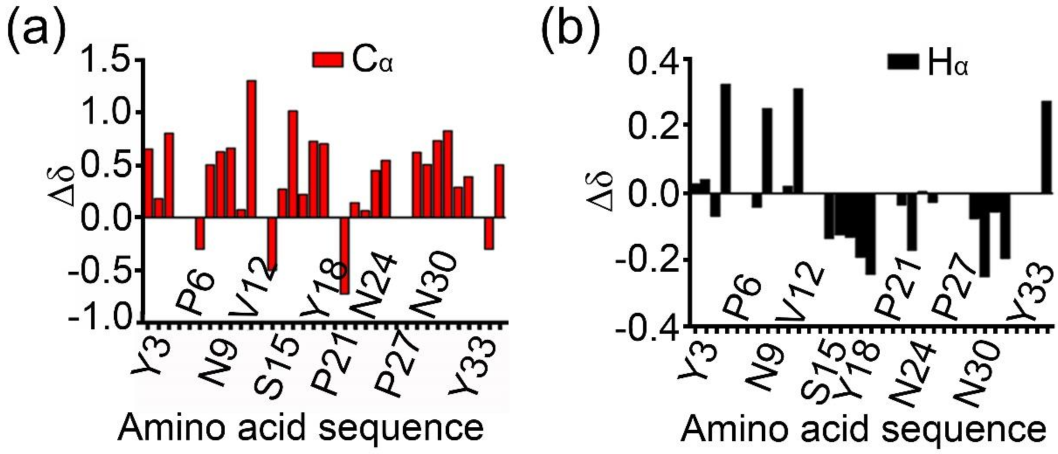


**Supplementary figure 15.** (a) Chemical shift deviations (Cα) plot of NS36 (b) Chemical shift deviations (Hα) plot of NS36.

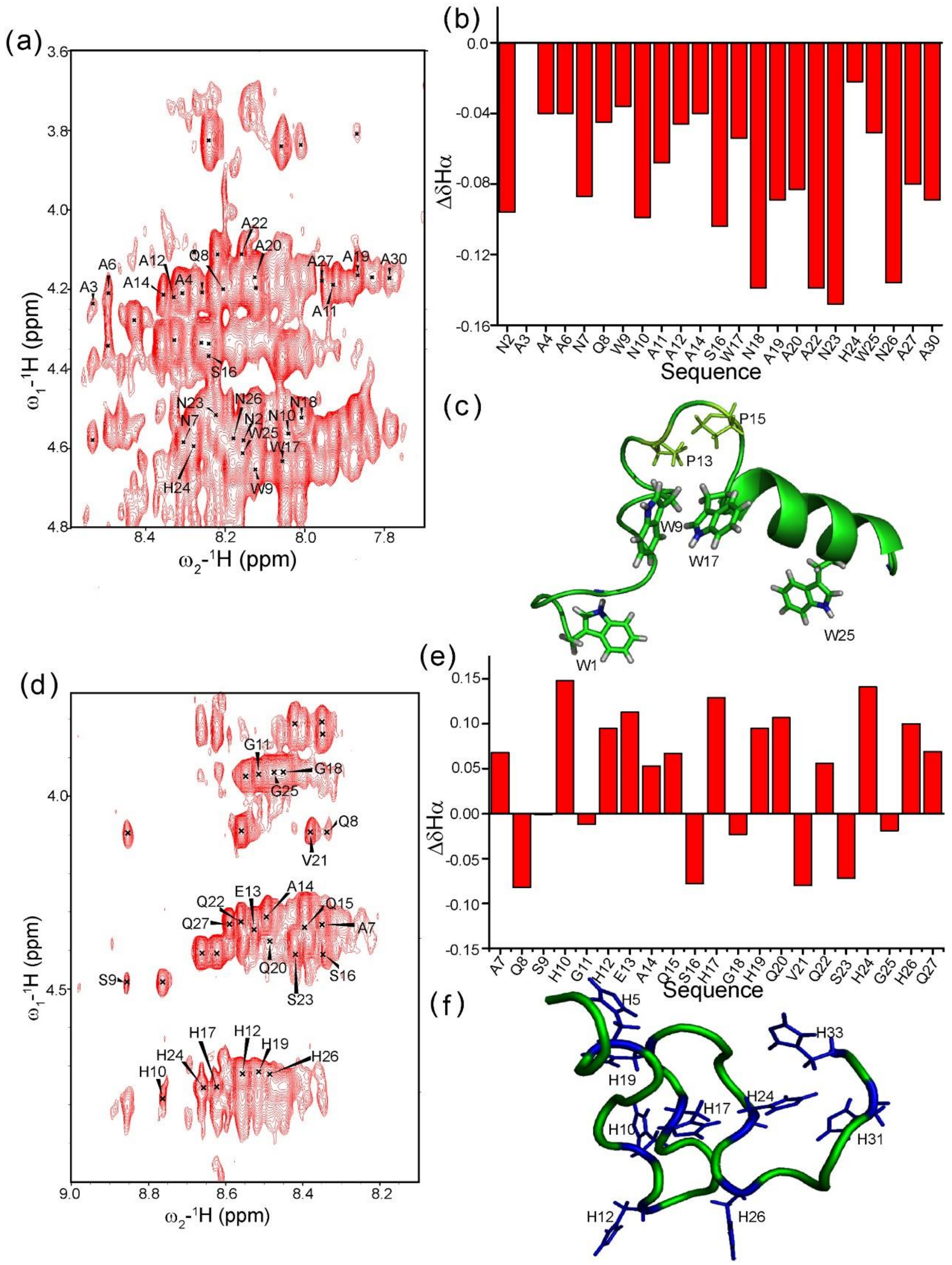


**Supplementary figure 16.** (a) 2D $^1$H-$^1$H NOESY spectrum of WA30 peptide displaying Hα region and noesy connectivities (only Hα spins of WA30 are marked for clarity) (b) Chemical shift deviations (Hα) plot of WA30 (c) Representative energy minimized structure of WA30 showing π-π interactions of W9-W17 while the other side chains of Trp (W1 and W25) remain exposed. (d) $^1$H-$^1$H NOESY spectrum of QH33 peptide displaying Hα region and noesy connectivities (only Hα spins of QH33 are marked for clarity) (e) Chemical shift deviations (Hα) plot of QH33 (f) Representative energy minimized structure of QH33 with sidechains of His marked in blue.

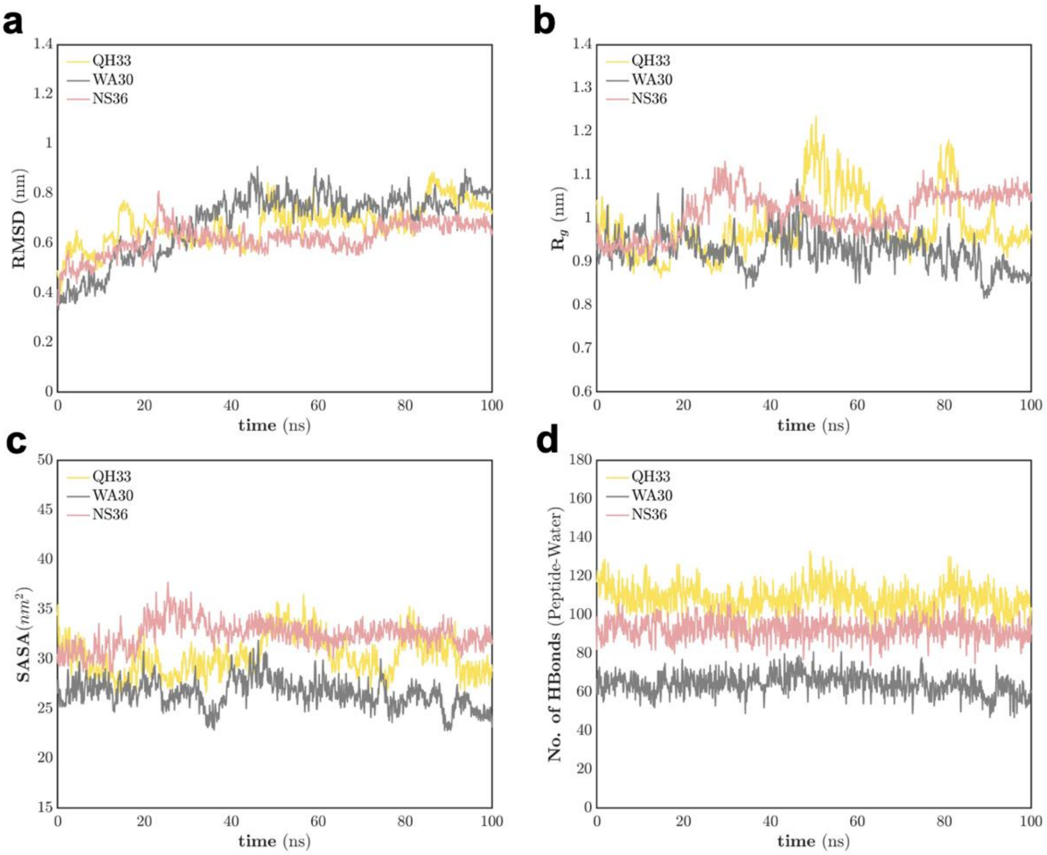


**Supplementary figure 17.** (a) Evolution of the root-mean-square deviation (RMSD) of simulated ICPs in water relative to the chosen NMR structure as the reference. (b) Evolution of the radius of gyration (Rg) of simulated ICPs, representing the compactness of the structures. (c) Evolution of the solvent accessible surface areas (SASA) of simulated ICPs, indicating the exposure of the molecules to the surrounding solvent. (d) Evolution of the number of hydrogen bonds formed between ICPs and water molecules during the simulations.

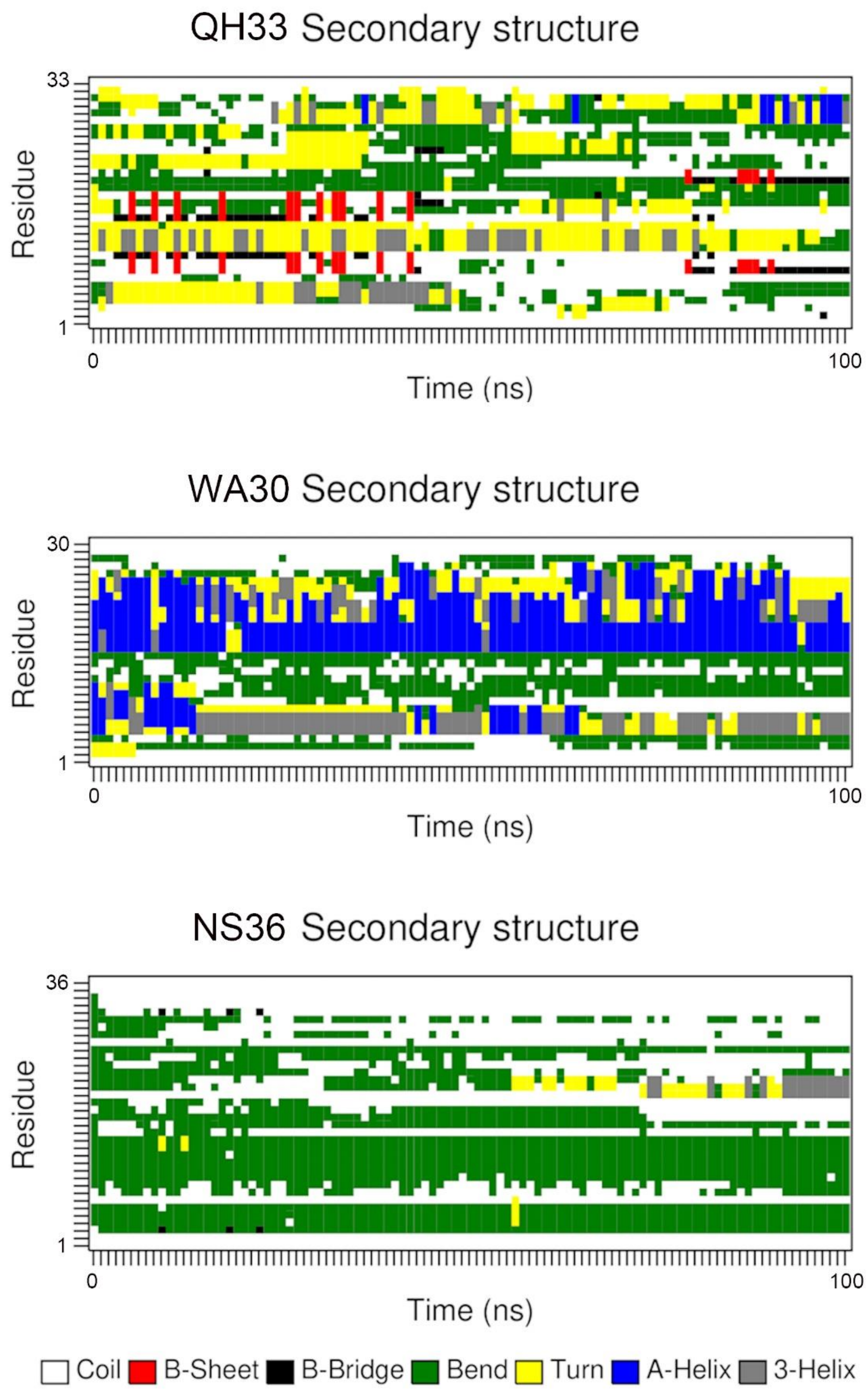


**Supplementary figure 18.** Temporal evolution of ICP secondary structures during aqueous MD simulations. The corresponding color codes representing each secondary structure are depicted in the legend bar at the bottom of the figure.

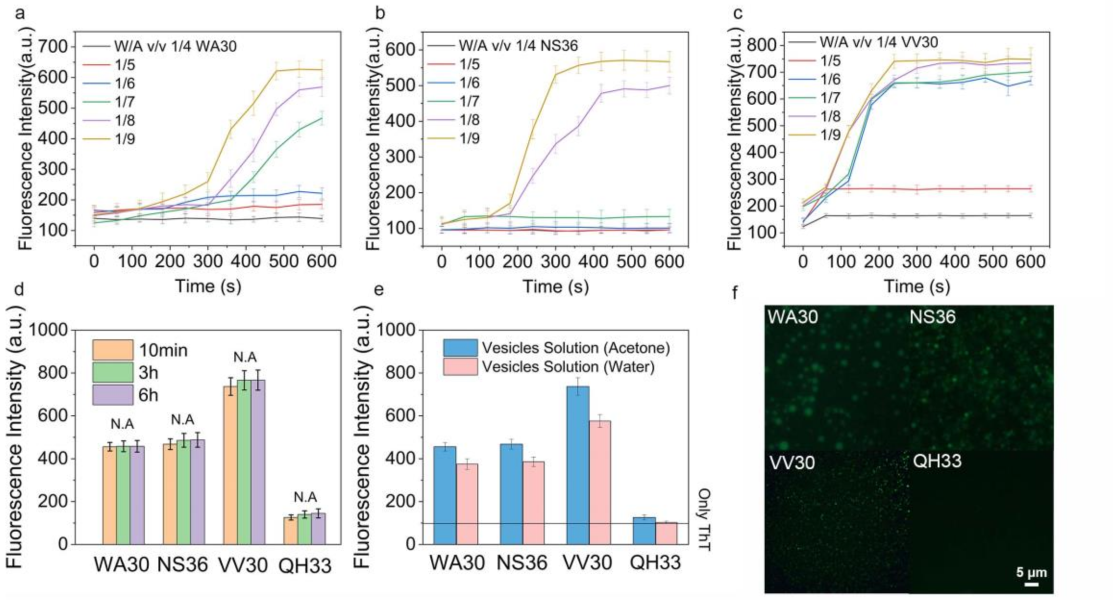


**Supplementary figure 19.** The kinetic analysis of ICPs WA30(a), NS36(b) and VV30(c) transformation structure information monitored though ThT staining in gradiometric composed mixed solvents. (d) Fluorescence spectra of ICPs transformation structure information monitored though ThT staining after vesicles formation within different time scale. (e) Structural changes of crosslinked vesicles after re-dialysis into the aqueous environment. (f) Fluorescence optical micrograph of peptide vesicles staining with 0.5 mg $mL^{-1}$ ThT. Scale bar is 5 μm.

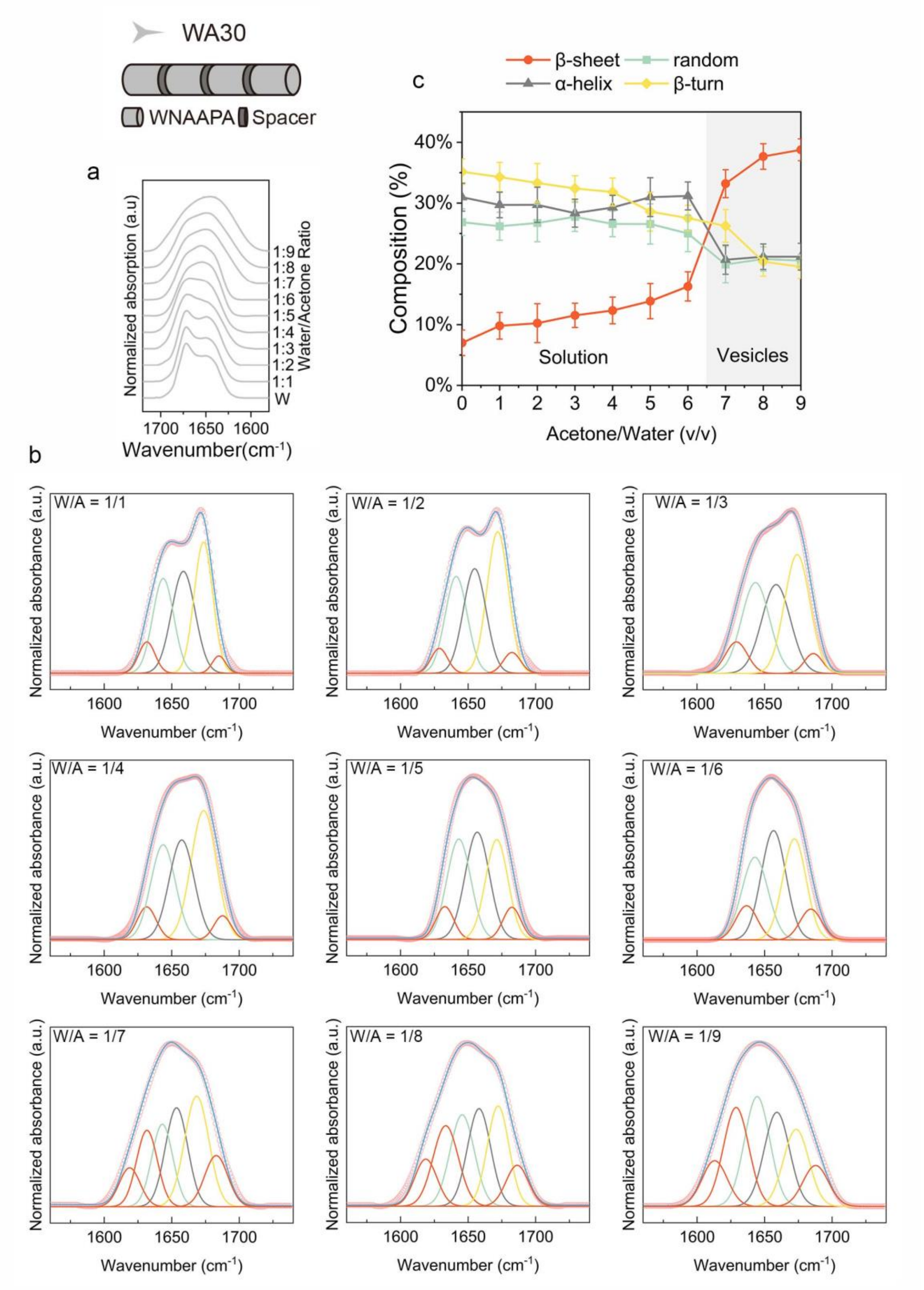


**Supplementary figure 20.** (a) The amide I ATR-FTIR spectra of WA30 under different water to acetone ratio, data are representative from n =3 independent assays. (b) FTIR deconvolution analysis of the amide I of WA30 under different water to acetone ratio. (c) The proportion of WA30 secondary structure based on FTIR deconvolution fitting; data are means ± SD from n = 3 independent fitting.

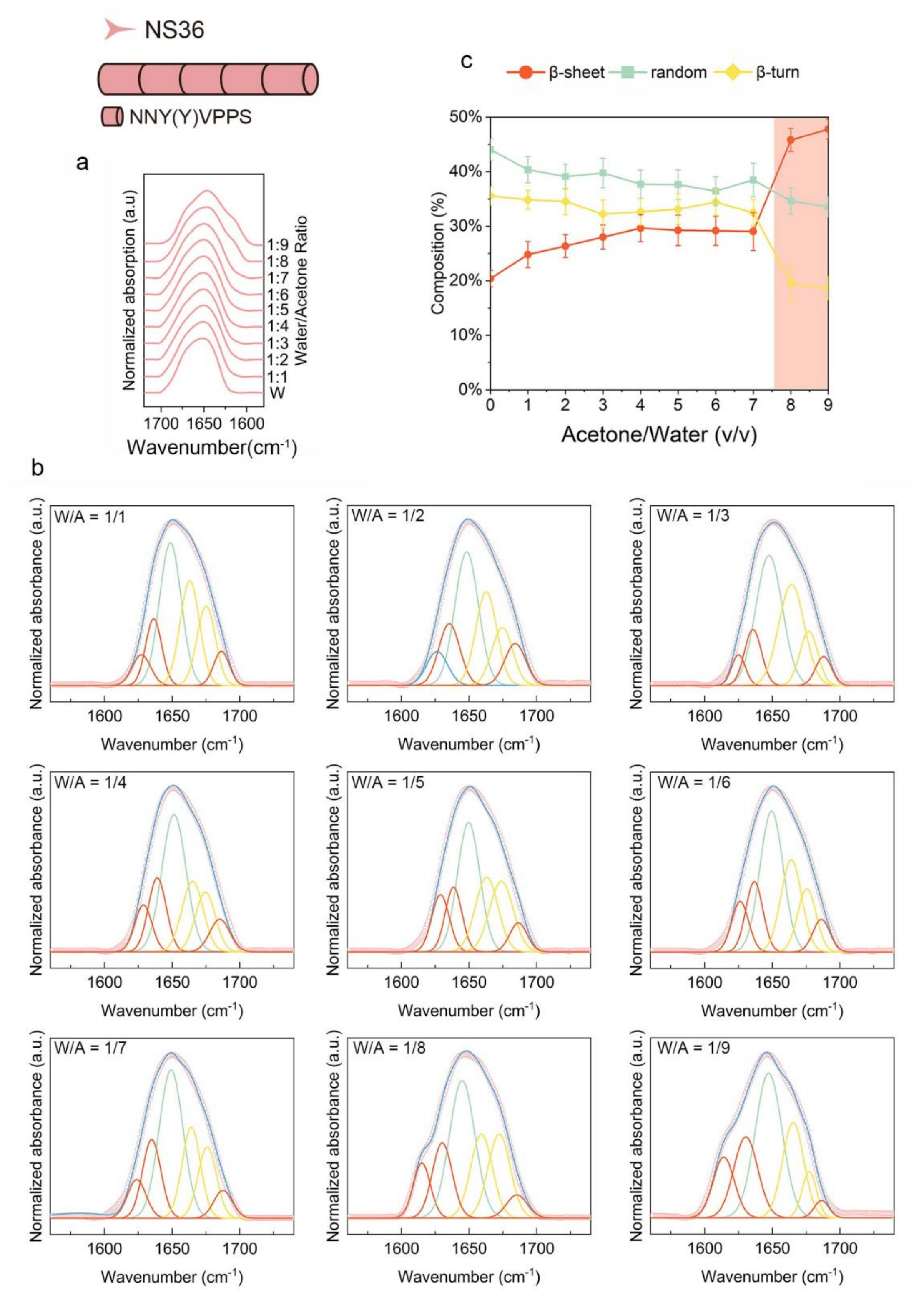


**Supplementary figure 21.** (a) The amide I ATR-FTIR spectra of NS36 under different water to acetone ratio, data are representative from n =3 independent assays. (b) FTIR deconvolution analysis of the amide I of NS36 under different water to acetone ratio. (c) The proportion of NS36 secondary structure based on FTIR deconvolution fitting; data are means ± SD from n = 3 independent fitting.

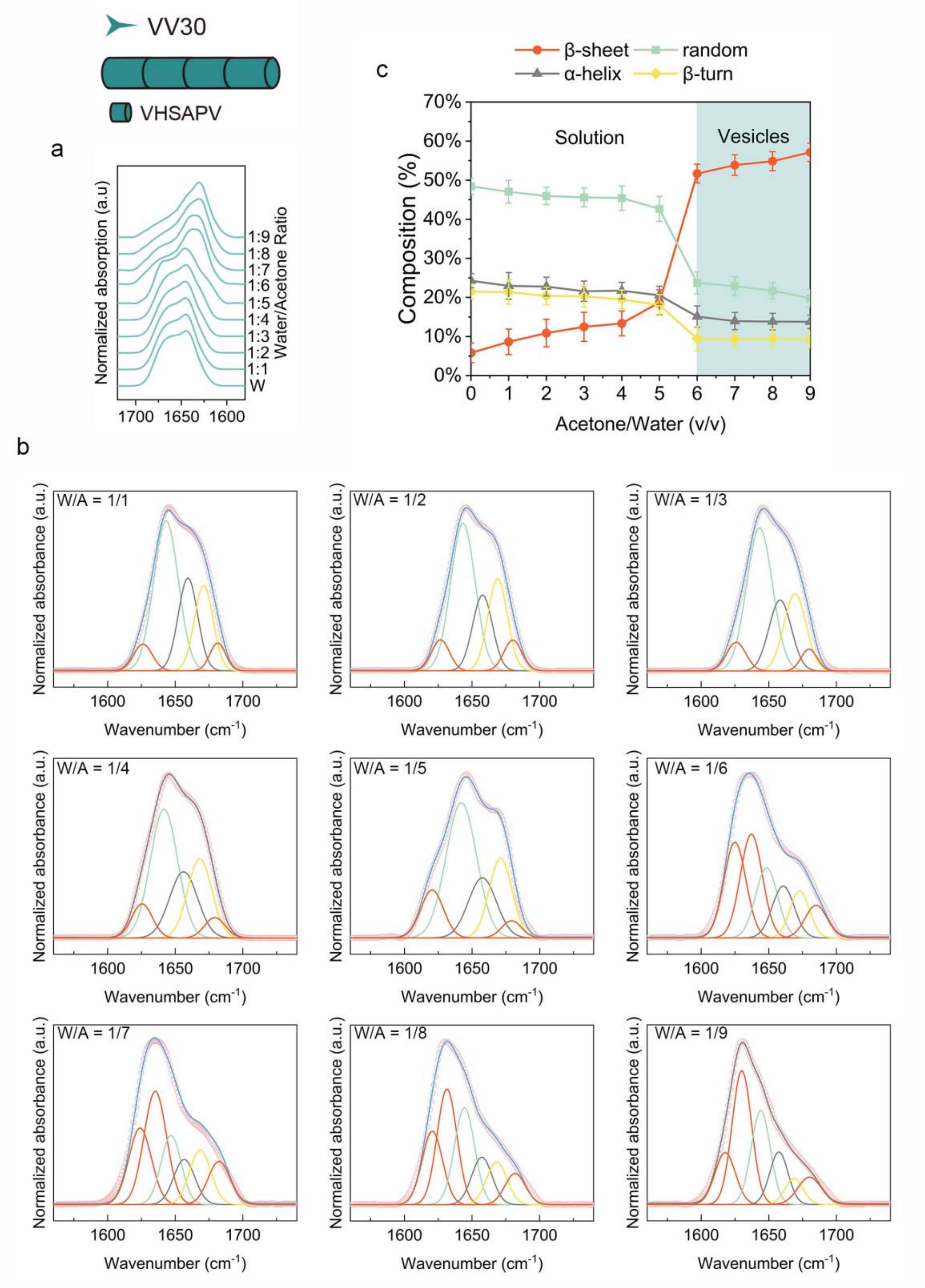


**Supplementary figure 22.** (a) The amide I ATR-FTIR spectra of VV30 under different water to acetone ratio, data are representative from n =3 independent assays. (b) FTIR deconvolution analysis of the amide I of VV30 under different water to acetone ratio. (c) The proportion of VV30 secondary structure based on FTIR deconvolution fitting; data are means ± SD from n = 3 independent fitting.

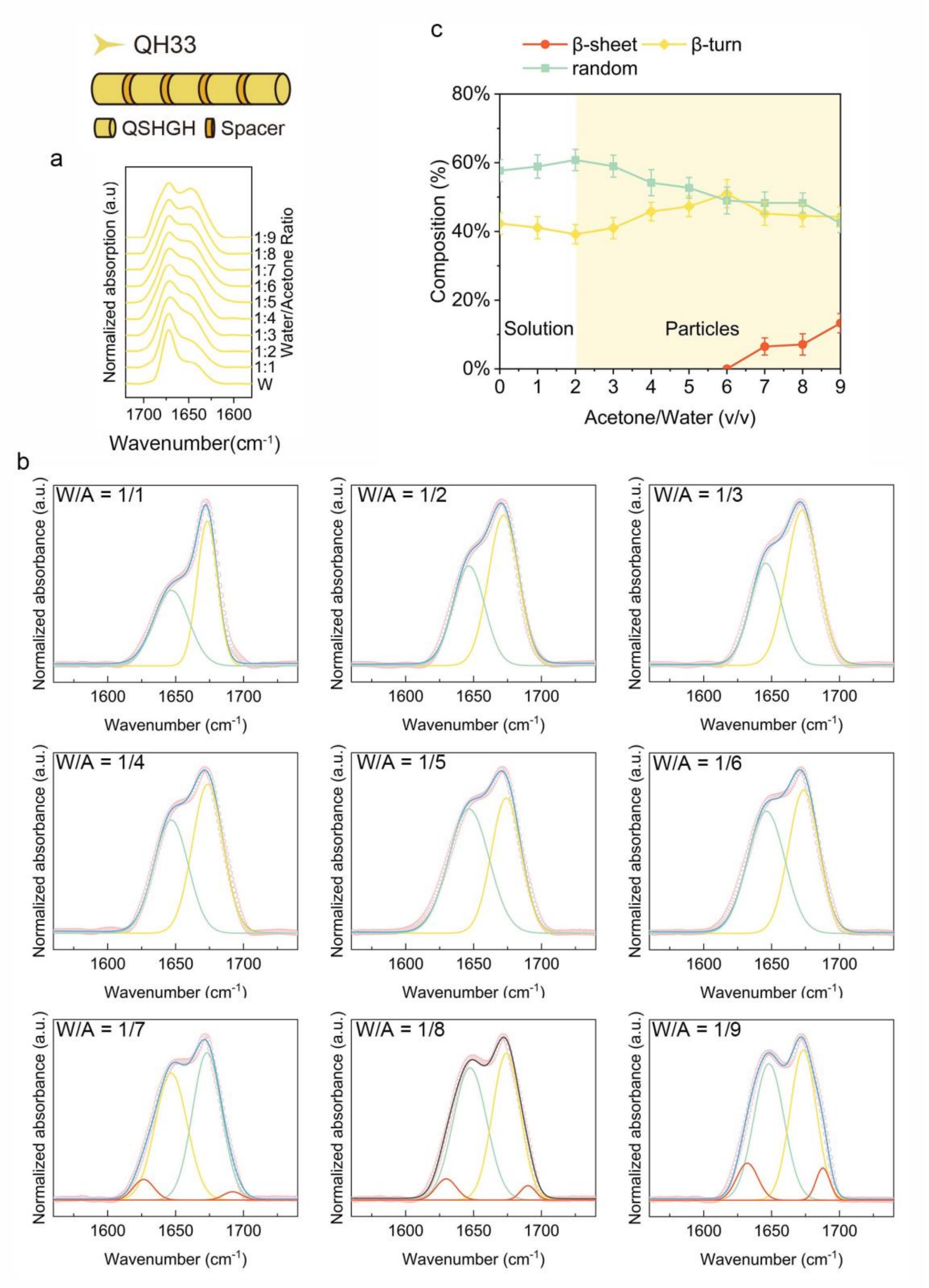


**Supplementary figure 23.** (a) The amide I ATR-FTIR spectra of QH33 under different water to acetone ratio, data are representative from n =3 independent assays. (b) FTIR deconvolution analysis of the amide I of QH33 under different water to acetone ratio. (c) The proportion of QH33 secondary structure based on FTIR deconvolution fitting; data are means ± SD from n = 3 independent fitting.

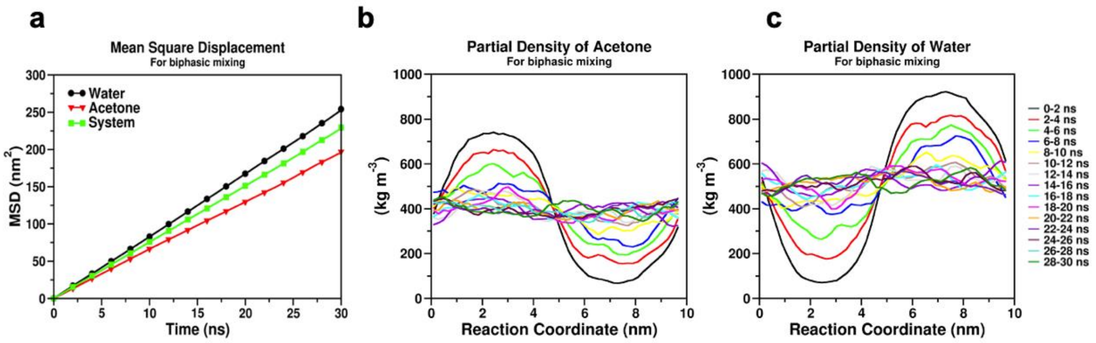


**Supplementary figure 24.** (a) Evolution of mean square displacement (MSD) for water, acetone, and the system. (b) Partial density plot showing the distribution of acetone along the reaction coordinate with 2 ns intervals. (c) Partial density plot showing the distribution of water along the reaction coordinate with 2 ns intervals.

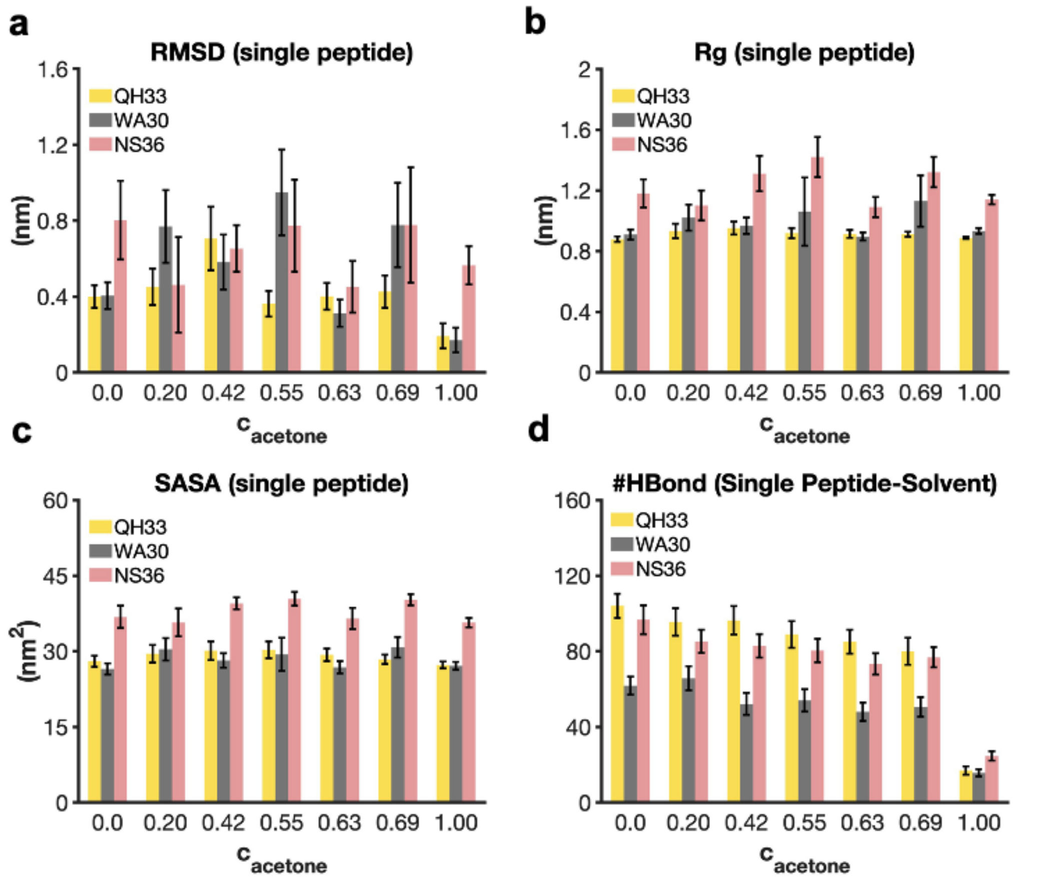


**Supplementary figure 25.** (a) Root-mean-square deviation (RMSD) of a peptide as a function of acetone concentration. (b) Radius of gyration (Rg) of a peptide as a function of acetone concentration. (c) Solvent accessible surface area (SASA) of a peptide as a function of acetone concentration. (d) Number of hydrogen bonds between a peptide and solvent molecules (water and/or acetone) as a function of acetone concentration. Data represent means ± standard deviation (SD) obtained from simulation trajectories.

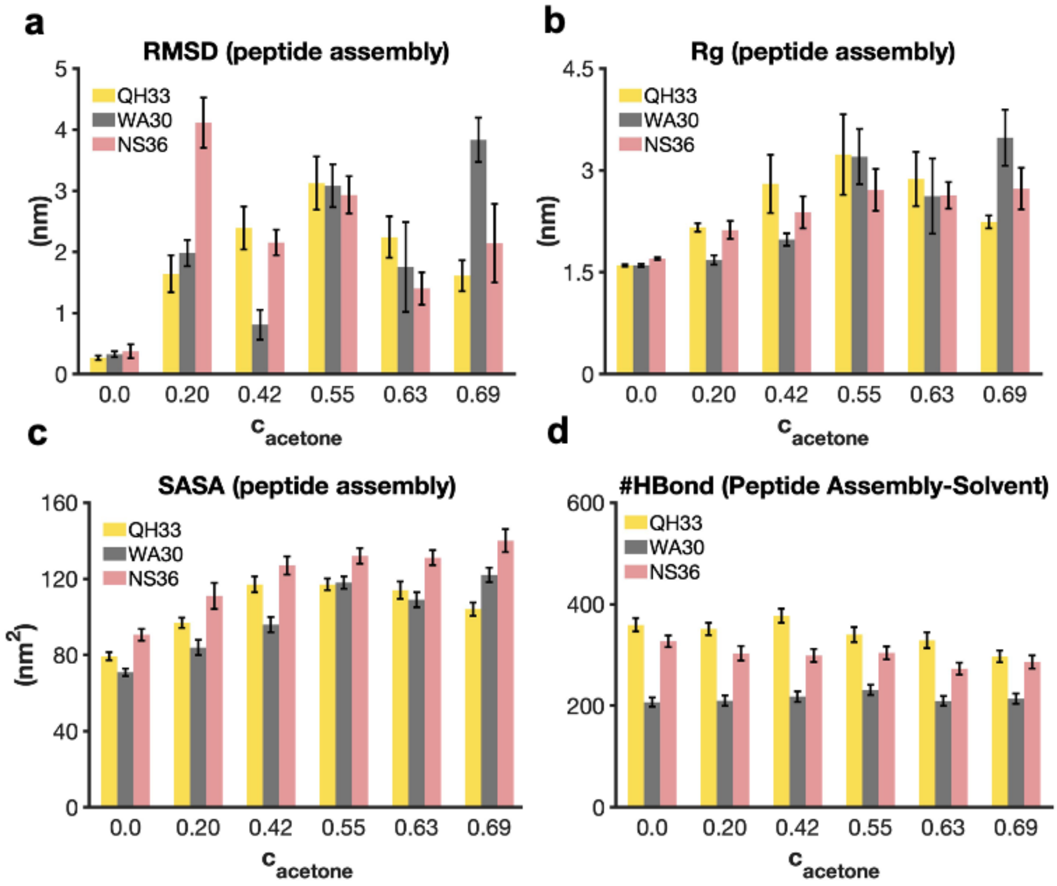


**Supplementary figure 26.** (a) Root-mean-square deviation (RMSD) of the peptide assembly as a function of acetone concentration. (b) Radius of gyration (Rg) of the peptide assembly as a function of acetone concentration. (c) Solvent accessible surface area (SASA) of the peptide assembly as a function of acetone concentration. (d) Number of hydrogen bonds between the peptide assembly and solvent molecules (water and/or acetone) as a function of acetone concentration. Data represent means ± standard deviation (SD) obtained from simulation trajectories.

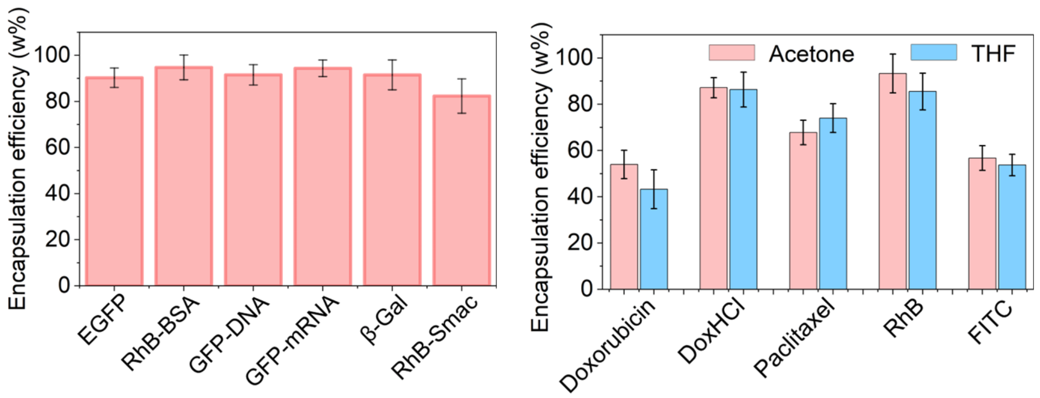


**Supplementary figure 27.** Recruitment efficiency of biomacromolecules(a) and small molecules(b) by NS36 capsules and the mean ± SD (column with error bar) of n = 3 independent measurements.

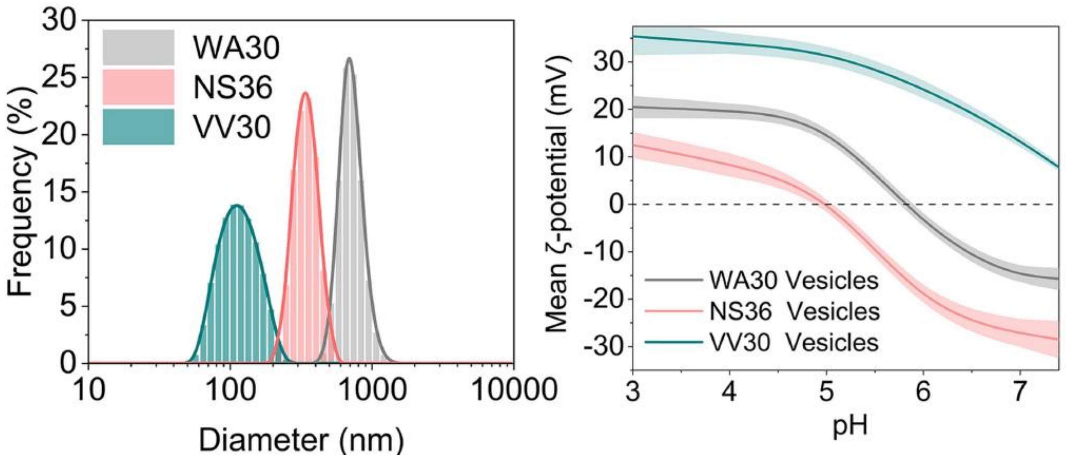


**Supplementary figure 28.** (a) Size distributions of ICPs capsules in PBS; (b) zeta potentials of ICPs capsules in different pH buffer and the mean ± SD of n = 3 independent measurements.

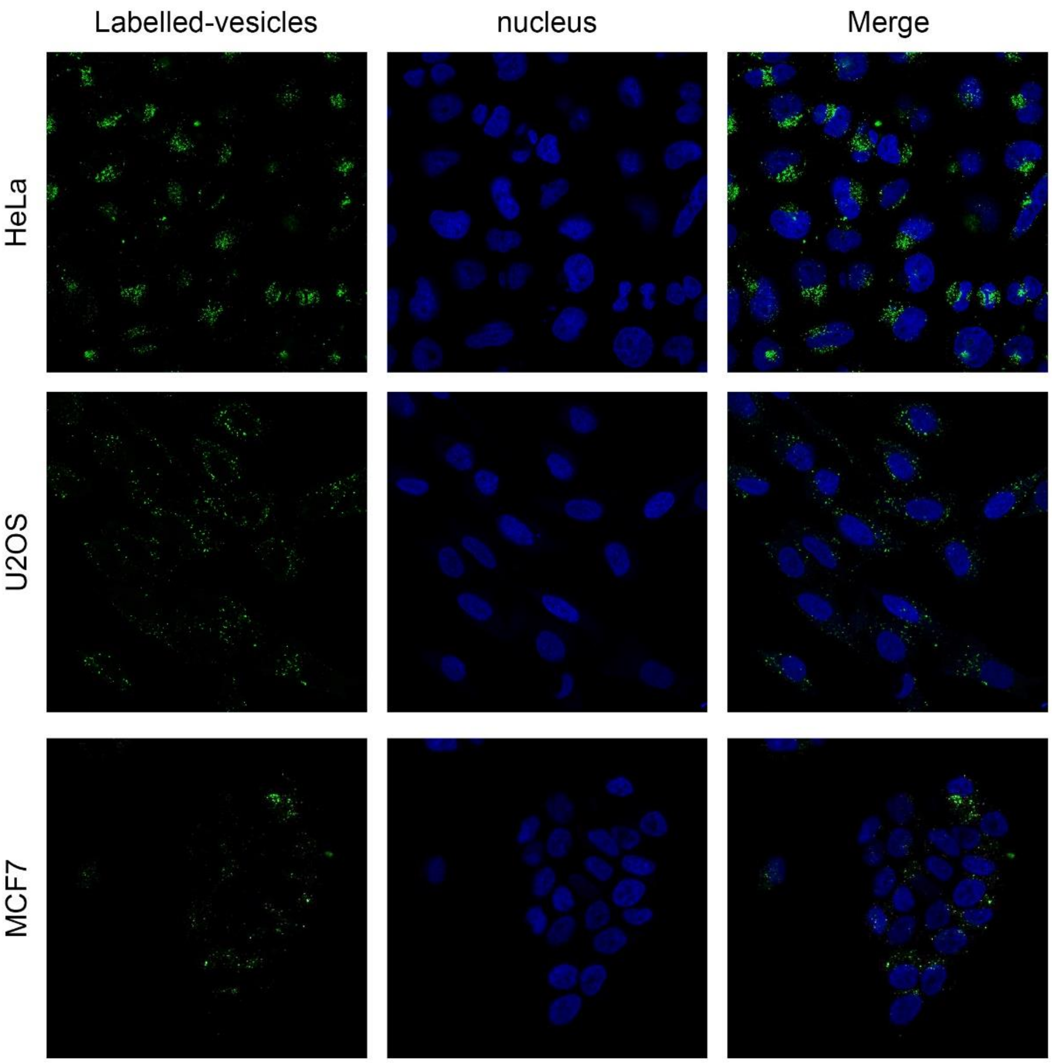


**Supplementary figure 29.** Confocal microscopy images of different cell line internalization of the NS36 nano-capsules.

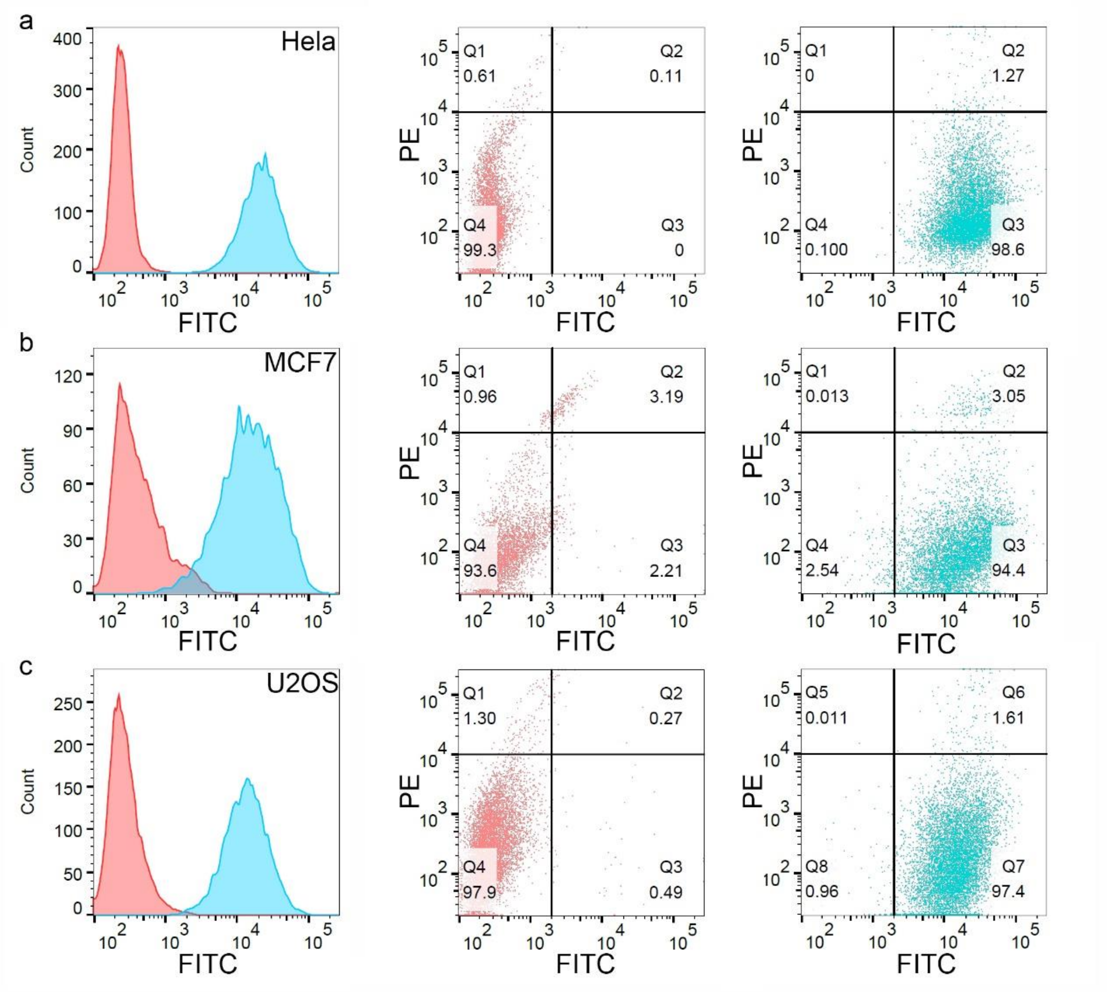


**Supplementary figure 30.** The FACS results of Different cell line internalization of the NS36 nano-capsules. The Cells with the PE intensity greater than $10^4$ were considered dead.

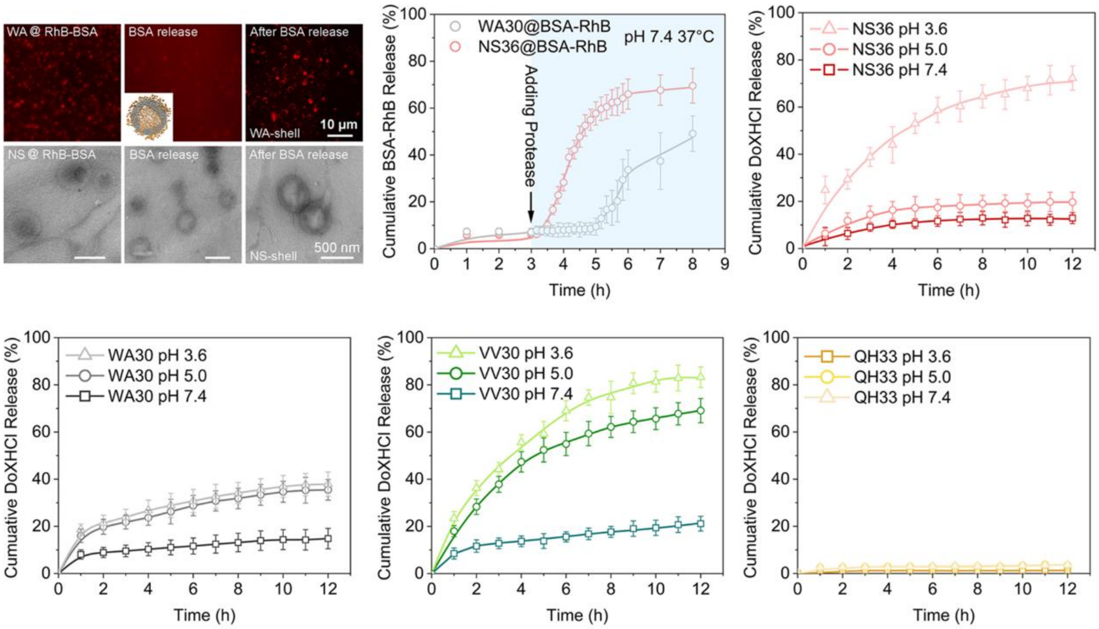


**Supplementary figure 31.** (a) Fluorescence microscopy and TEM images of the release of RhB labeled BSA loaded on NS36 and WA30 capsules; (b) breaking the capsules structure by adding 500 ng $mL^{-1}$ enzymes in vitro to release internal macromolecular cargos and the mean ± SD (error bar) of n = 3 independent measurements; (c) DoXHCl(10 μg $mL^{-1}$) releasing dynamics of NS36(c), WA30(d), VV30(e) and QH33(f) capsules under different pH, the mean ± SD of n = 3 independent measurements.

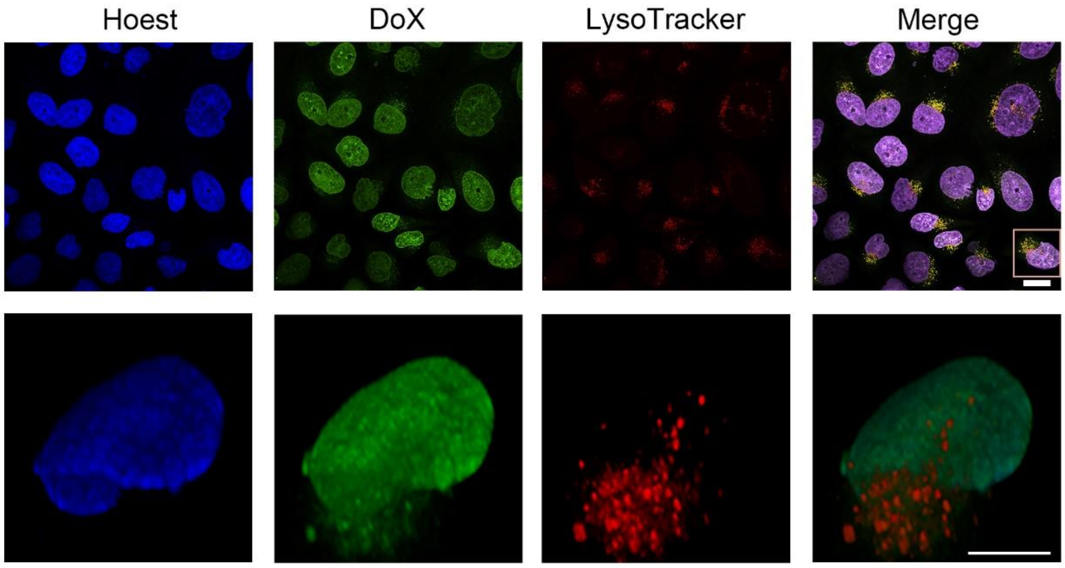


**Supplementary figure 32.** Confocal microscopy images of Hela cells treated with FITC-labeled BSA and DoXHCl co-loaded NS36 vesicles for 24 h. The delivery result was confirmed by n = 3 independent assays. The DoXHCl was showed by green, and the lysosome was stained by lysoTracker. The DoXHCl were generally co-localized with lysosome and nucleus. Scale bar for original image is 20 μm, and for partial enlarged 3D reconstructed image is 10 μm. The partially enlarged 3D reconstructed image showed the release of DoX from the lysosome which was co-located with nano-capsules.

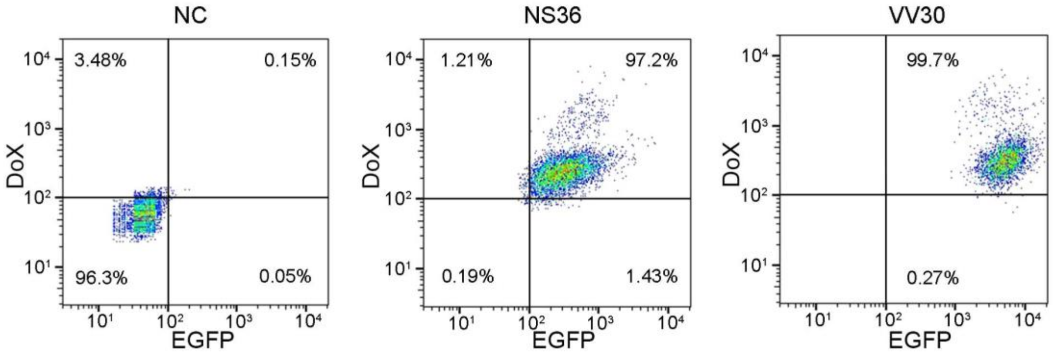


**Supplementary figure 33.** The FACS analysis of nano-capsules co-delivery of DoX and EFGP to HeLa cell for 4h. After four hours of internalizing the NS36 and VV30 nano-capsules, the two experimental groups showed similar DoX intensity, while EGFP intensity showed significant differences. We believe that the phenomenon was also related to the proton sponge effect, where the remaining NS36 nano-capsules in lysosome were stayed in the lower pH environment. This low pH environment will cause fluorescence quenching of EGFP and result in lower signal intensity, meanwhile the EGFP loaded in VV30 which escaped into the cytoplasm did not have this concern.

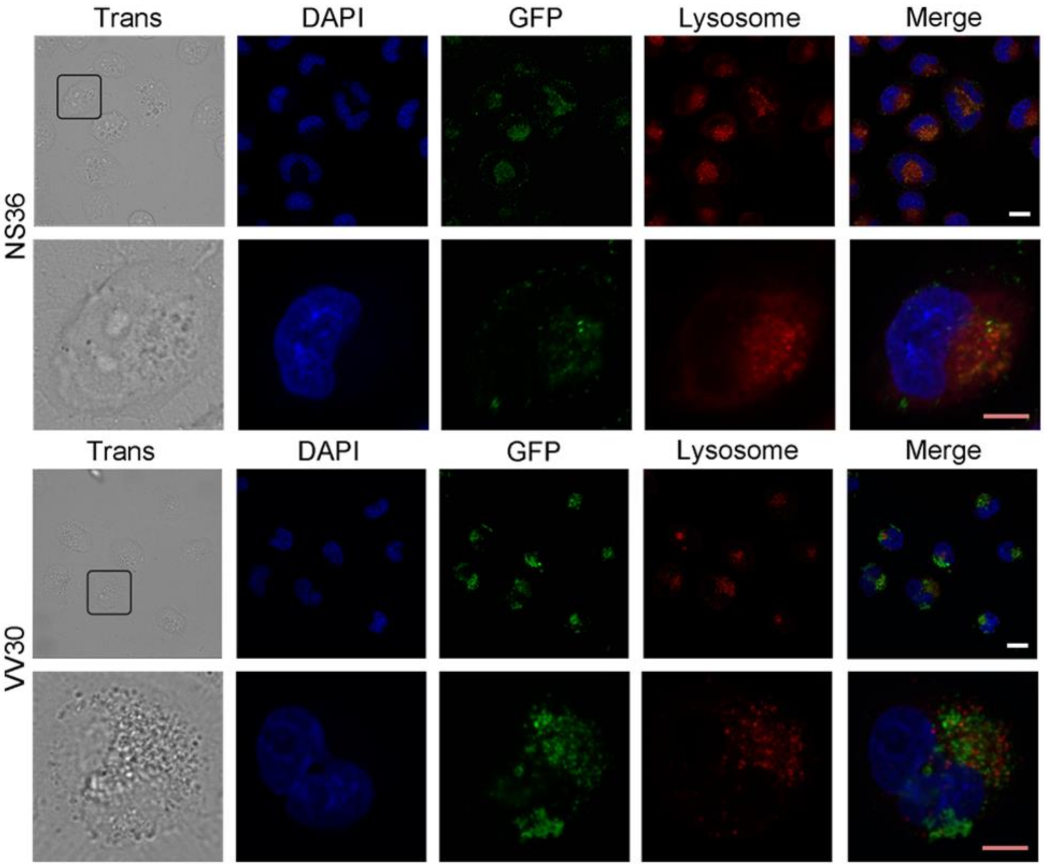


**Supplementary figure 34. Cell internalization of different peptide nano-capsules.** The NS36 nano-capsules entered lysosome after being internalized by HeLa cells, and confocal fluorescence images showed high fluorescence co-localization of lysosome and NS36 nano-capsules. However, the vesicles of VV30 were almost not co-localized with lysosomes after internalization. We believe that the reason for this difference was due to the proton sponge effect caused by the His group on the surface of VV30 NS36 nano-capsules, which also caused the VV30 NS36 nano-capsules to escape into the cytoplasm. The white scale bar is 20 μm, the pink scale bar is 10 μm.

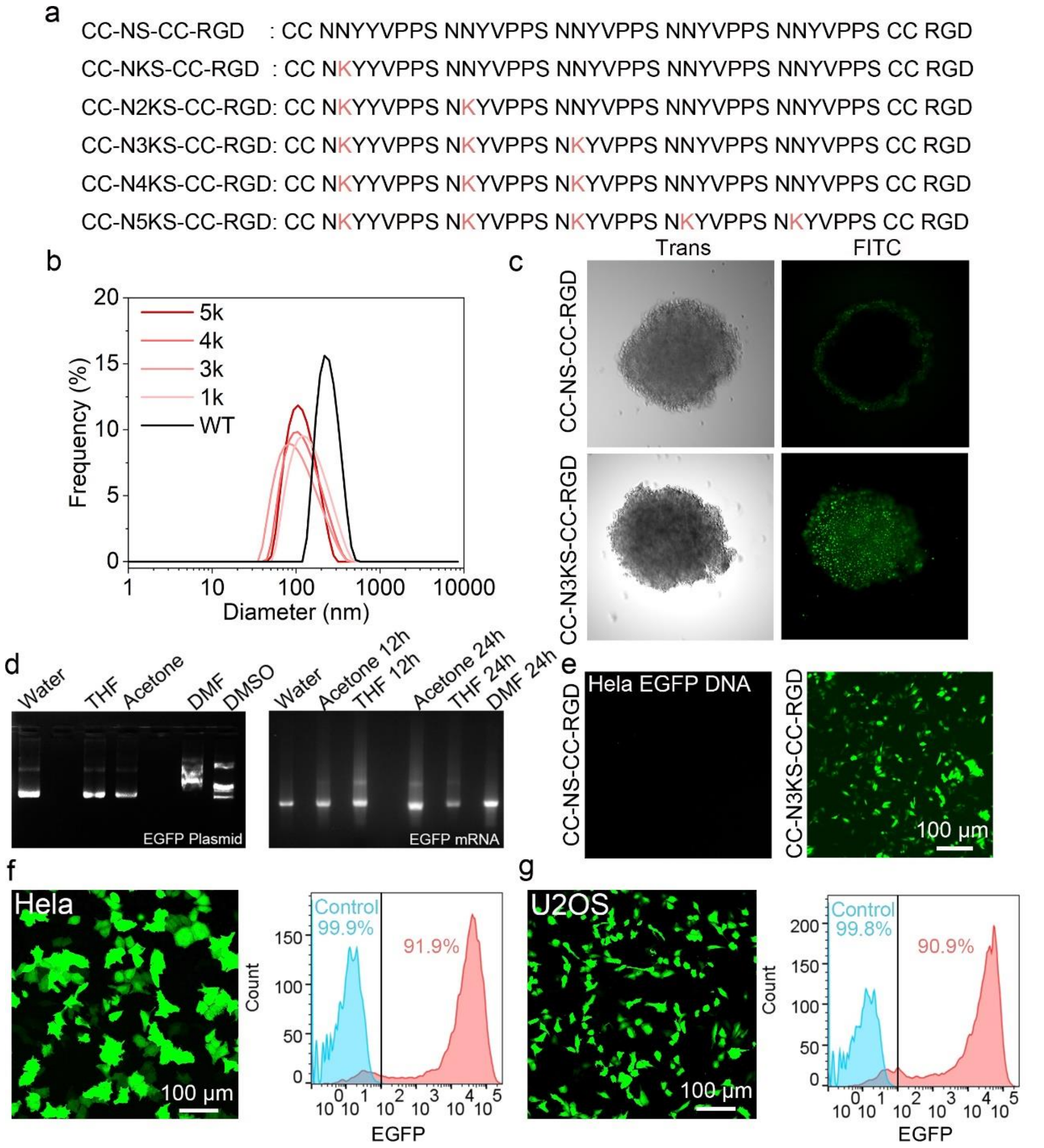


**Supplementary figure 35. Intracellular plasmid and mRNA transfection though NS36 mutations nano-capsules.** (a)The sequence of designed NS36 mutations which suitable for plasmid and mRNA delivery; (b) Size distributions of NS36 mutations capsules in PBS; (c) the internalization of NS36 mutations capsules by a HeLa cell mass; (d) the effect of different organic solvents on the stability of EGFP nucleic acid; (e) confocal microscopy images and FACS analysis of HeLa cells treated with EGFP plasmid loaded CC-N3KS-CC-RGD capsules; (f) confocal microscopy images and FACS analysis of HeLa cells treated with EGFP mRNA loaded CC-N3KS-CC-RGD capsules; (g) confocal microscopy images and FACS analysis of U2OS cells treated with EGFP mRNA loaded CC-N3KS-CC-RGD capsules.

**Supplementary Information Reference**


1 Martin, M. Cutadapt removes adapter sequences from high-throughput sequencing reads. *EMBnet. journal* **17**, 10-12 (2011).

2 Haas, B. J. *et al.* De novo transcript sequence reconstruction from RNA-seq using the Trinity platform for reference generation and analysis. *Nature protocols* **8**, 1494-1512 (2013).

3 Guo, Q. *et al.* Hydrogen-bonds mediate liquid-liquid phase separation of mussel derived adhesive peptides. *Nature Communications* **13**, 5771 (2022).

4 Hornak, V. *et al.* Comparison of multiple Amber force fields and development of improved protein backbone parameters. *Proteins: Structure, Function, and Bioinformatics* **65**, 712-725 (2006).

5 Jorgensen, W. L., Chandrasekhar, J., Madura, J. D., Impey, R. W. & Klein, M. L. Comparison of simple potential functions for simulating liquid water. *The Journal of chemical physics* **79**, 926-935 (1983).

6 Wang, J., Wolf, R. M., Caldwell, J. W., Kollman, P. A. & Case, D. A. Development and testing of a general amber force field. *Journal of computational chemistry* **25**, 1157-1174 (2004).

7 Caleman, C. *et al.* Force field benchmark of organic liquids: density, enthalpy of vaporization, heat capacities, surface tension, isothermal compressibility, volumetric expansion coefficient, and dielectric constant. *Journal of chemical theory and computation* **8**, 61-74 (2012).

8 Hess, B., Bekker, H., Berendsen, H. J. & Fraaije, J. G. LINCS: A linear constraint solver for molecular simulations. *Journal of computational chemistry* **18**, 1463-1472 (1997).

9 Hoover, W. G. Canonical dynamics: Equilibrium phase-space distributions. *Physical review A* **31**, 1695 (1985).

10 Parrinello, M. & Rahman, A. Polymorphic transitions in single crystals: A new molecular dynamics method. *Journal of Applied physics* **52**, 7182-7190 (1981).

11 Qian, X., Lucherelli, M. A., Corcelle, C., Bianco, A. & Gao, H. Mechanics of biosurfactant aided liquid phase exfoliation of 2D materials. *Forces in Mechanics* **8**, 100098 (2022).

12 Klimovich, P. V., Shirts, M. R. & Mobley, D. L. Guidelines for the analysis of free energy calculations. *Journal of computer-aided molecular design* **29**, 397-411 (2015).
